# SynapseNet: Deep Learning for Automatic Synapse Reconstruction

**DOI:** 10.1101/2024.12.02.626387

**Authors:** Sarah Muth, Frederieke Moschref, Luca Freckmann, Sophia Mutschall, Ines Hojas-Garcia-Plaza, Julius N. Bahr, Arsen Petrovic, Thanh Thao Do, Valentin Schwarze, Anwai Archit, Kirsten Weyand, Susann Michanski, Lydia Maus, Cordelia Imig, Nils Brose, Carolin Wichmann, Ruben Fernandez-Busnadiego, Tobias Moser, Silvio O. Rizzoli, Benjamin H. Cooper, Constantin Pape

## Abstract

Electron microscopy is an important technique for the study of synaptic morphology and its relation to synaptic function. The data analysis for this task requires the segmentation of the relevant synaptic structures, such as synaptic vesicles, active zones, mitochondria, presynaptic densities, synaptic ribbons, and synaptic compartments. Previous studies were predominantly based on manual segmentation, which is very time-consuming and prevented the systematic analysis of large datasets. Here, we introduce SynapseNet, a tool for the automatic segmentation and analysis of synapses in electron micrographs. It can reliably segment synaptic vesicles and other synaptic structures in a wide range of electron microscopy approaches, thanks to a large annotated dataset, which we assembled, and domain adaptation functionality we developed. We demonstrated its capability for (semi-)automatic biological analysis in two applications and made it available as an easy-to-use tool to enable novel data-driven insights into synapse organization and function.

## Introduction

Analyzing electron micrographs of synapses is a key technique for understanding synaptic morphology and its connection to function and plasticity. To date, this task is predominantly performed manually^1,2^ and constitutes a bottleneck in the study of synaptic ultrastructure. The last decade has seen the adoption of deep learning to automate such analysis tasks, largely through methods that build on the UNet architecture^3,4^, resulting in tools for cell segmentation in light microscopy^5–7^ or organelle segmentation in electron microscopy7,8. However, to our knowledge, an equivalent tool for the analysis of synapses in electron microscopy is still missing due to the lack of training data and a robust segmentation method. Here, we introduce SynapseNet, which closes this gap. SynapseNet segments vesicles and other synaptic structures in different kinds of electron micrographs, enabling (semi-)automatic reconstruction and analysis of synapses. We created a large annotated dataset of synaptic structures in electron tomography, which we used to train deep neural networks for different segmentation tasks. We also developed an algorithm for domain adaptation that enables applying these networks to different imaging conditions and we showed that SynapseNet can speed up relevant analysis tasks. It is available as a python library and graphical user interface.

Electron microscopy serves as a particularly useful technique for resolving the subcellular organization of inter-neuronal synapses and establishing corresponding links to defined functional states^1,9–12^. Corresponding studies are based either on serial section transmission electron microscopy^13–15^, room temperature electron tomography^12,16–18^ or scanning electron microscopy^19–21^. Cryogenic electron tomography^22–24^ can also reveal molecular information. Quantitative analysis of these data requires the segmentation of subcellular compartments (i.e. presynaptic terminal, postsynaptic compartment), synaptic organelles (e.g. synaptic vesicles, mitochondria, smooth endoplasmic reticulum), and subsynaptic compartments (e.g. active zone, postsynaptic density). Based on these segmentations, researchers can analyze vesicle pools and distances between structures, or derive other measurements to identify correlates of functional synapse states. The majority of previous studies relied on manual segmentation with tools such as IMOD^25^, RECONSTRUCT^26^ or SynasEM^2^. However, such manual segmentation is very time consuming, impeding systematic analyses of large datasets. Automation of synapse reconstruction would thus mark a significant advancement for the field. Prior work has addressed some of the segmentation tasks: Methods for vesicle segmentation have been proposed for electron tomography based on classical image analysis^27^ and deep learning^28^, for cryogenic electron tomography based on template matching^23,29^ and deep learning^30,31^, and for transmission electron microscopy based on deep learning^32,33^. For mitochondrion segmentation robust deep learning based tools exist^7,8^, but there are no methods for other synaptic structures of major functional relevance, such as the active zone. MemBrain^34^ and TomoSegMemTV^35^ provide robust segmentation of plasma membranes in cryogenic electron tomography, and Dragonfly^36^ offers functionality for data annotation and training of deep neural networks, but they do not specifically target the synapse. None of this prior work offers a unified solution to the reconstruction of synapses, as the methods either target only a specific structure (synaptic vesicle, mitochondrion or membrane) or require user annotations to train a network for each structure. Furthermore, the vesicle segmentation methods only work for specific electron microscopy sample-preparation techniques (e.g. aldehyde fixation, cryo fixation) and/or imaging modalities (e.g. room-temperature vs. cryogenic electron tomography), but do not generalize to other conditions. See <u>Materials and Methods</u> for a more detailed discussion of other tools.

Our tool, SynapseNet, provides a comprehensive solution for segmentation and analysis of synapses in electron micrographs; Figure 1 gives an overview. Our main contributions are:

1. Assembling a large dataset of electron tomograms with over 117,000 annotated synaptic vesicles and annotations for active zones, including those of ribbon synapses, mitochondria, and synaptic compartments.
2. Training deep neural networks for segmentation based on this dataset. We trained networks for 2D and 3D segmentation of synaptic vesicles as well as 3D segmentation networks for the other structures.
3. Introducing a domain adaptation method that improves these networks when applied to different conditions, e.g. different sample preparation techniques or different imaging modalities. This method does not require any additional annotations, broadening the applicability of SynapseNet.
4. Implementing functionality for common analysis tasks, including vesicle pool assignment and distance measurements, that can be used via SynapseNet’s graphical user interface or its python library.

**Figure 1:**
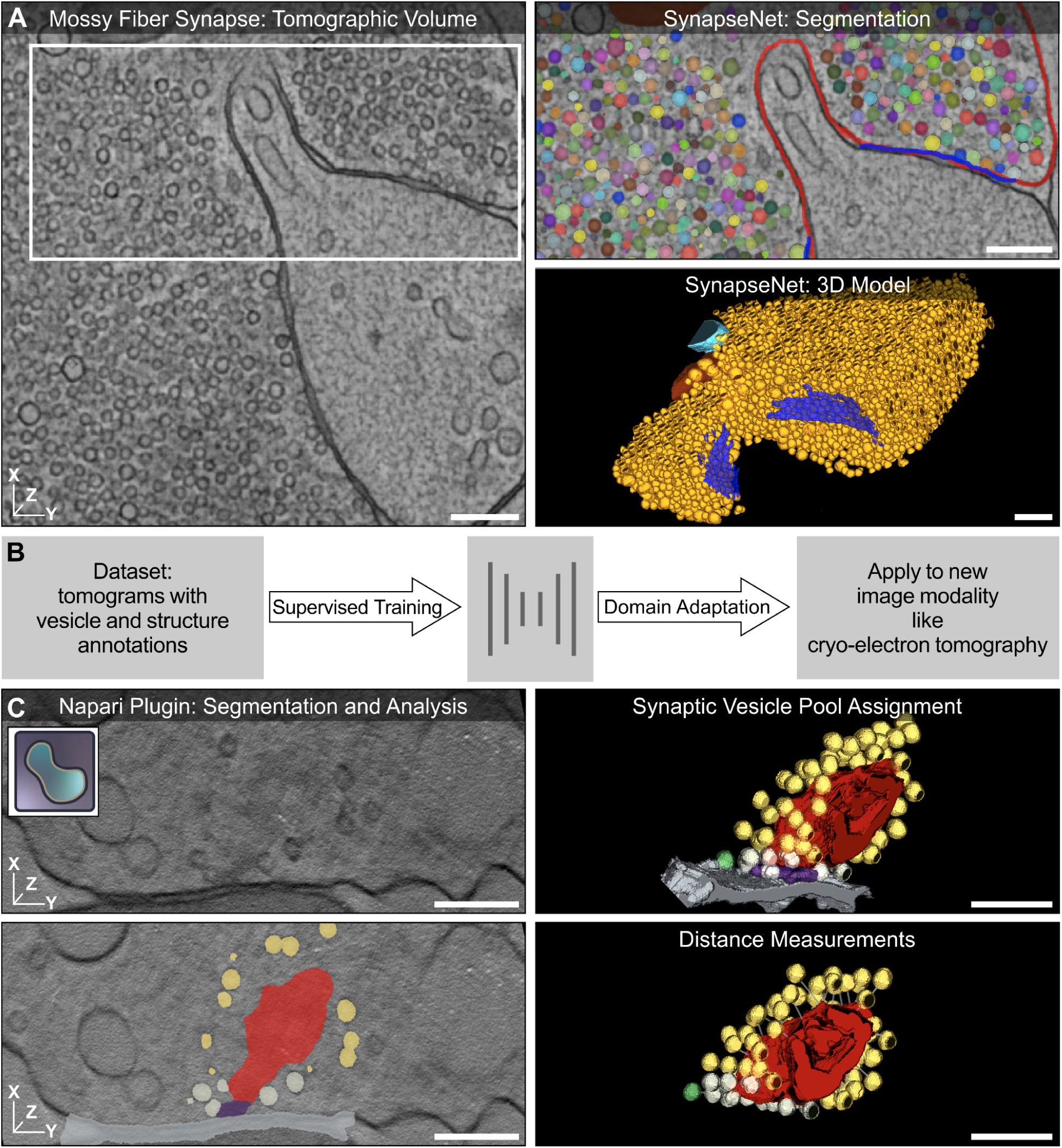
Overview of SynapseNet, a tool for automatic synapse reconstruction in electron micrographs. **A.** Virtual section of an electron tomogram of a mossy fiber synapse with synaptic structures segmented by SynapseNet shown in the zoom-in and the 3D rendering: synaptic vesicles (different colors in zoom-in, orange in 3D), mitochondria (red and cyan), and active zones (blue). See Supplementary Figure 1 for further examples. **B.** Schematic overview of SynapseNet’s segmentation functionality. We trained deep neural networks to segment synaptic structures on annotated data (left) and developed a domain adaptation algorithm (right) to adapt these networks to new imaging modalities without the need for additional annotations. **C.** Applications of SynapseNet. We implemented a python library and a graphical user interface (napari^37^ plugin) to perform segmentation (bottom left) and analysis tasks, such as vesicle pool assignment (top right) and distance measurement (bottom right). The example shows an inner ear ribbon synapse with segmentation of synaptic vesicles, divided into three different pools, (yellow, gray, green), the ribbon (red), the presynaptic density (purple), and the active zone membrane (gray). The scale bars represent 200 nm.

SynapseNet is already in use for multiple on-going studies of synaptic ultrastructure. Here, we demonstrated how it can be used for analyzing synaptic morphology in hippocampal Schaffer collateral and inner ear ribbon synapses. We expect that SynapseNet will speed up the analysis of synaptic ultrastructure by circumventing manual segmentation, thus enabling new insights into synapse biology by a comprehensive ultrastructure reconstruction that would not be possible otherwise.

## Results

Here, we present (i) the evaluation of the segmentation networks underlying SynapseNet, (ii) the evaluation of the domain adaptation functionality that enables applying it to different imaging conditions, (iii) two example analyses that can be performed with it, and (iv) the tools we provide to make SynapseNet available to researchers.

### Automatic segmentation in electron tomography

Training deep neural networks for segmentation tasks requires a large dataset with annotations. To our knowledge, such a dataset was not previously available for the segmentation of synaptic ultrastructure. We created it by assembling data from previous studies that had employed electron tomography for analyses of synapses^11,12,16,17,38,39^ and by annotating previously unpublished electron tomograms. The resulting dataset contains annotations for synaptic vesicles, active zones, mitochondria, ribbon synapse structures, and synaptic compartments. Figure 2A and 2B show an overview of the number of tomograms and annotated objects. A detailed description of all underlying data is given in <u>Sample Preparation and Data Acquisition</u> and the data annotation procedure is described in <u>Data Annotation</u>.

**Figure 2:**
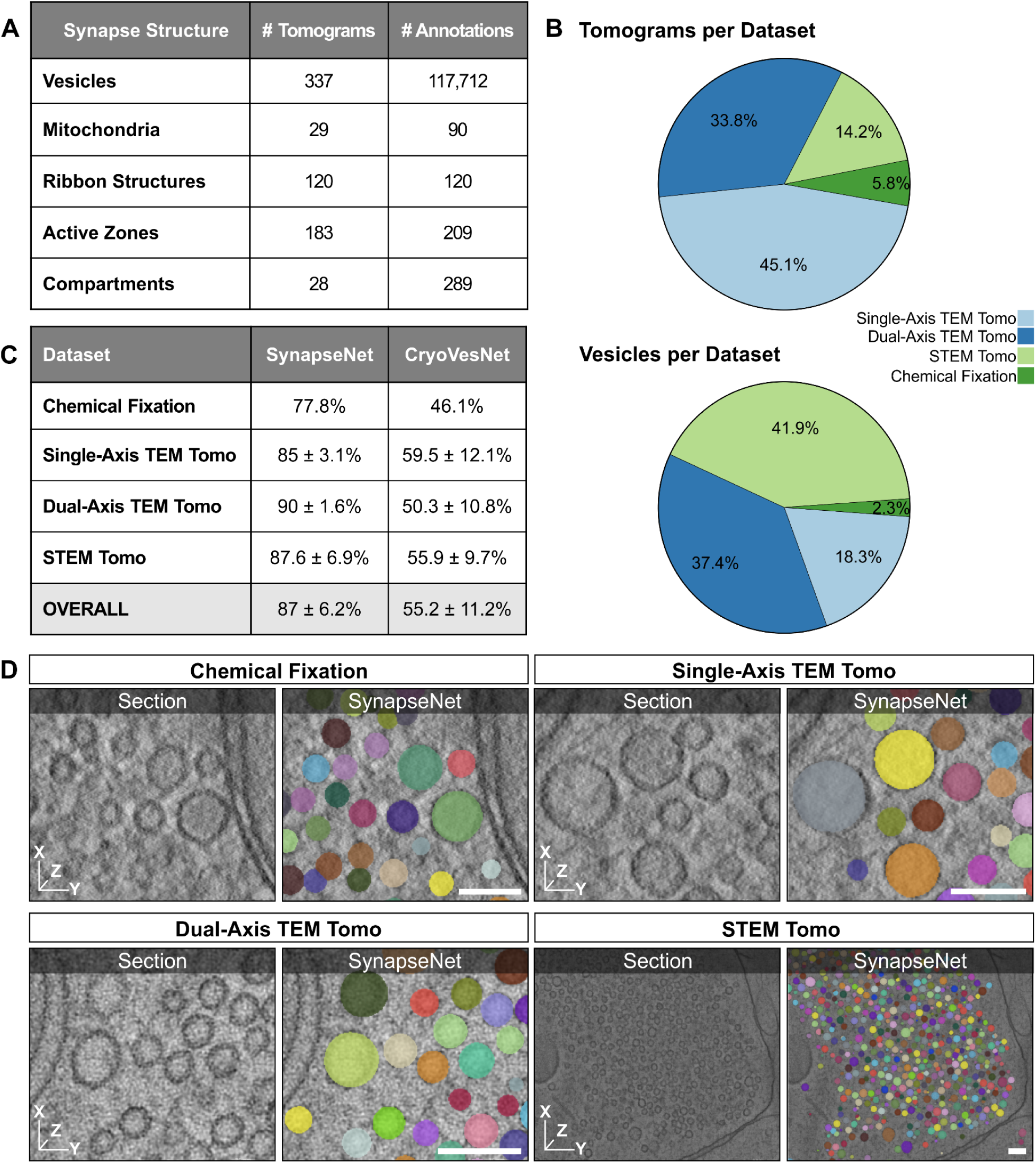
Training data and vesicle segmentation. **A.** Overview of the training data for the different synaptic structures in our dataset. **B.** Fraction of tomograms (top) and annotated vesicles (bottom) distributed over the four different imaging conditions in the dataset. Please refer to <u>Sample Preparation and Data Acquisition</u> for an explanation of these conditions. **C.** Evaluation of the vesicle segmentation results, using the F1-Score at an intersection over union of 50%, of SynapseNet and CryoVesNet^30^. We evaluated the segmentations on a separate test split and differentiate between the four different image conditions. **D.** Qualitative segmentation results from SynapseNet. We show individual virtual sections (left) and the segmentation result overlay (right). Each individual vesicle identified by our method is drawn with a different color. The scale bars represent 100 nm.

Based on these data, we trained multiple neural networks for different segmentation tasks:

- A network for synaptic vesicle segmentation in volumetric data using a 3D UNet^4^.
- A network for synaptic vesicle segmentation in 2D image data using a 2D UNet^3^.
- A network for active zone segmentation in volumetric data using a 3D UNet.
- A network for mitochondrion segmentation in volumetric data using a 3D UNet.
- A network for ribbon, presynaptic density and active zone membrane segmentation in ribbon synapses, in volumetric data using a 3D UNet.
- A network for synaptic compartment segmentation in volumetric data using a 3D UNet.

All networks make use of a similar architecture and training hyperparameters, see <u>Supervised</u> <u>segmentation</u> for details. Our main focus was on synaptic vesicle segmentation, which is the most time consuming step in manual data analysis. We evaluated the network for vesicle segmentation in volumetric data on a separate split of the dataset with annotations. We compared the segmentation results to the annotations using the F1-Score at an intersection over union of 50% (see <u>Segmentation metrics</u>). Figure 2C presents the corresponding results, including a comparison to CryoVesNet^30^, the only other easy-to-use tool for volumetric vesicle segmentation that we are aware of. Figure 2D shows qualitative segmentation results, separately for four different types of electron tomography in the dataset. Our method yielded segmentations of high quality with agreement of segmentations and annotations between 77.8% and 90%, depending on the tomogram type, and an overall agreement of 87%. It outperformed CryoVesNet, which was only trained on cryogenic electron tomography and thus performed poorly for room-temperature electron tomography. The results for the 2D vesicle segmentation network are shown in Supplementary Figure 2A. Here, the same data split was used as in the training for the 3D model, however, the network was trained and evaluated on individual virtual sections of the tomograms. Compared to the 3D segmentation network, it achieved a lower segmentation quality of overall 70.7%. This is due to the fact that it cannot make use of 3D context. The 2D evaluation was also affected by the ambiguity of annotations in 2D, where the exact beginning and end of a vesicle across the depth axis is often ill-defined due to the missing wedge effect in tomography. Nevertheless, providing a network for 2D vesicle segmentation is important to support analysis of synapses in serial section transmission electron microscopy, which is a common approach in the field. Our domain adaptation experiments, see the next section, demonstrated that the 2D network is up to this task.

We also evaluated the active zone segmentations on a separate test set, measuring the segmentation quality using the Dice coefficient. Active zones are well separated spatially, so this metric is appropriate here. The evaluation results are listed in Supplementary Figure 2B; SynapseNet reached an overall score of 54.90%. Visual evaluation showed that all active zones were found, but they were often smaller than the annotations or erroneously picked up parts of the postsynaptic membrane. For the ribbon synapses we evaluated the segmentations on 12 separate test tomograms. We segmented two different structures: the ribbon, an electron dense structure that holds vesicles close to the active zone, and the presynaptic density, which can be identified by its higher electron density in these data. Here, we used the Dice coefficient to compare segmentations and annotations because ribbon and presynaptic density are well separated spatially, so distinguishing ribbons / presynaptic densities from each other is not an issue. We measured an agreement of 85.78 +-6.68% for the ribbon segmentation and 34.63 +-29.41% for the presynaptic density segmentation. I.e., the ribbon was reliably segmented, but the presynaptic density was not. The ribbon is a large structure that can be identified based on its high electron density and the network can reliably find and delineate it. Synaptic ribbons can have a complex morphology, explaining the remaining error in the segmentation. In contrast, the presynaptic density is a small structure of lower electron density and thus smaller contrast. Hence, the network only correctly identifies it in about a third of cases. Supplementary Figure 3 shows example segmentation results for the ribbon synapse structures. For mitochondria and synaptic compartments we had fewer annotated tomograms, as such we evaluated the corresponding segmentations only qualitatively. We performed visual inspection of the segmentation results on 35 STEM tomograms that were not used for training. We found that 159 out of 166 mitochondria (95.78 %) were correctly segmented. In three cases two mitochondria were wrongly merged into a single object in the segmentation and in one case a mitochondrion was split up into multiple pieces. In addition, 22 objects were wrongly identified as mitochondria, usually large endo- or exosomes. For synaptic compartments, we only checked the segmentations of presynaptic terminals. The most common error was compartments split into multiple pieces and we found an average of 1.31 +-0.75 pieces for the 74 terminals we checked, with a maximum of 5 pieces per compartment. We also qualitatively evaluated the active zone segmentation, as this imaging modality was not part of the test set used for quantitative evaluation in Supplementary Figure 2B. Here, we found that 78 of 116 active zones (67.24 %) were found, i.e. at least a sizable fraction of the active zone was predicted by SynapseNet. Example segmentations of mitochondria, active zones, and a synaptic compartment are shown in Supplementary Figure 2C and D.

### Domain adaptation for robust synaptic structure segmentation

Deep neural networks trained with supervised learning often exhibit poor generalization to different data distributions, so-called data from a different domain. This limitation also affects our segmentation networks. The segmentation quality generally deteriorates when applied to different sample preparations or image modalities, for example, cryogenic electron tomography or serial section transmission electron microscopy. The field of domain adaptation studies techniques to improve the generalization to unseen data distributions. Here, we developed a new domain adaptation algorithm for 2D and 3D segmentation that is easy to apply in practice and that improves segmentation quality without requiring additional annotations. It uses a student-teacher approach^40,41^ to adapt the pretrained network to the new domain and is an extension of our previous work on domain adaptation for 2D segmentation^42^. See <u>Domain</u> <u>adaptation</u> for details.

To evaluate this method we applied it to different settings:

- Adapting 3D vesicle segmentation to different electron tomography data.
- Adapting 3D vesicle segmentation to cryogenic electron tomography.
- Adapting 2D vesicle segmentation to serial section transmission electron microscopy.

See <u>Sample Preparation and Data Acquisition</u> for details on the data. All domain adaptation experiments followed the same approach: The initial network was adapted to the new domain using our teacher-student adaptation method. This process was unsupervised, i.e. it did not require manual annotations in the new domain. We then evaluated the segmentation results on separate data from the new domain, with annotations, following the same evaluation procedure as before. The quantitative evaluation results are shown in Figure 3A and qualitative results in 3B. We compared to CryoVesNet for volumetric segmentation. For 2D segmentation we are not aware of another suitable method and thus did not compare to any other approach. The results showed an improved performance for all settings, ranging from modest improvements of 2-3% (2D TEM, EH), to clear improvements of 15-20% (IER, Cryo-64K). In the case of the cryogenic electron tomography datasets (Cryo-33K, Cryo-64K), CryoVesNet was better than our initial segmentation (before adaptation), however, SynapseNet performed on par with or better than CryoVesNet after adaptation. The 2D TEM dataset was challenging due to high vesicle density. Despite this, our network correctly identified ca. 75% of vesicles after adaptation, which would provide a good basis for semi-automatic analysis or further correction and model training in this setting. We also evaluated domain adaptation for vesicle segmentation in frog synapses^43^, which were imaged by serial section transmission electron microscopy. Here, our initial network found only very few vesicles due to the low contrast of these data. In this case, the segmentation quality deteriorated further after domain adaptation, thus showing that our domain adaptation only works if the initial predictions are of good-enough quality. Supplementary Figure 7B shows examples from these data.

**Figure 3:**
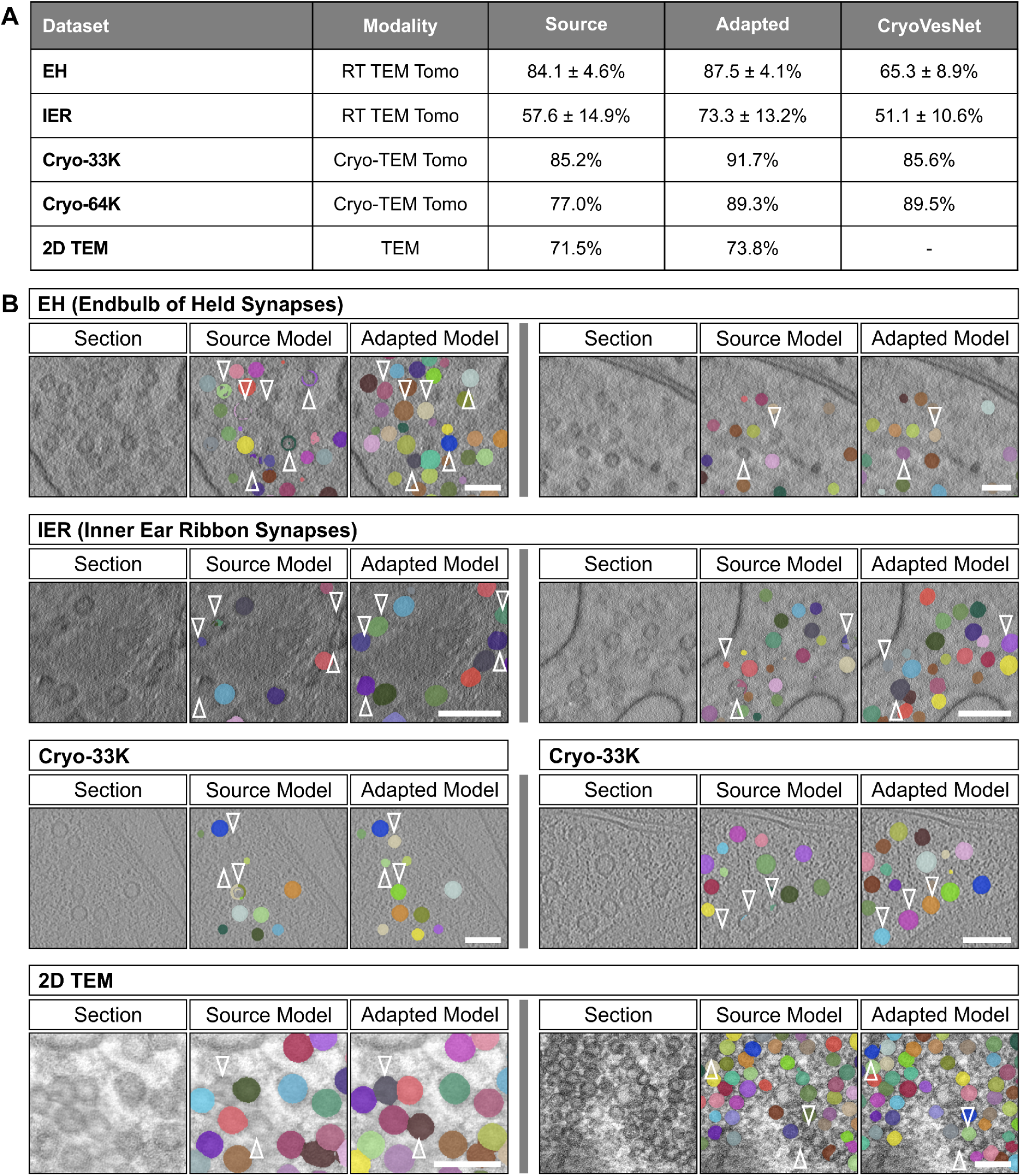
Domain adaptation for vesicle segmentation. **A.** Quantitative evaluation of different domain adaptation settings. We evaluated the results for four 3D datasets and a 2D dataset, (see <u>Sample Preparation and Data Acquisition</u> for details on the data). For each of them we report the metric for the model trained via supervised learning (‘Source’), for the model after domain adaptation (‘Adapted’), and for CryoVesNet (except for 2D data). The evaluation was performed on data with annotations, which was not used for the domain adaptation training. The evaluation procedure is the same as in Figure 2B. **B.** Example segmentations before and after adaptation. The arrows highlight vesicles for which the segmentation improved. The scale bars represent 100 nm.

### Automatic Reconstruction and Analysis

To demonstrate that SynapseNet can (semi-)automate the analysis of synaptic morphology, we performed biological evaluations for hippocampal Schaffer collateral and inner ear ribbon synapses. In the first analysis we compared automatic and manual annotations of synaptic vesicles and active zones to redo an already published analysis^17^ of Munc13-1/2^44^ and SNAP-25^45^ deficient synapses. We specifically selected these mutants for testing SynapseNet’s performance because in addition to functional deficits in synaptic signaling^44,46,47^, they exhibit reliable presynaptic ultrastructural phenotypes that manifest primarily at the level of synaptic vesicle organization and vesicle size (see Imig et al., 2014)^17^. Synapses deficient of Munc13 priming proteins or SNAP-25 SNARE complex components are characterized by an almost complete, or severe, loss of docked synaptic vesicles in contact with the active zone membrane, respectively. This observation supports the notion that morphologically docked synaptic vesicles serve as a reliable proxy for the molecularly-primed pool of fusion-competent vesicles. Electron tomography also resolved a statistically significant tendency for synaptic vesicles in Munc-13- and SNAP-25-deficient synapses to accumulate within 5-20 nm of the active zone. This latter observation provides morphological evidence in support of multiple, molecularly-regulated steps preceding synaptic vesicle priming and stimulus-evoked exocytosis. Finally, synaptic vesicles were larger in Munc13- and SNAP-25 deficient synapses compared to littermate controls. Although the mechanism(s) via which these mutations cause larger vesicles remain primarily speculative, the phenomenon is robust and has been observed in alternative preparations and imaging modalities^48^.

Here, we analyzed the tomograms from (Imig et al., 2014)^17^, manually segmenting synaptic vesicles and active zones for a subset of 20 tomograms (5 per knockout and respective control) with IMOD and automatically for all 101 tomograms, see <u>Synaptic Analysis</u> for the experimental set-up. We observed that only some of the dense core vesicles were correctly segmented by SynapseNet, likely due to the fact that they are underrepresented in the training data. We also observed that active zones were segmented completely in only 60% of the tomograms, and we corrected the active zone segmentations for the other 40%. The results for Munc13-½ double knock-out (DKO) and SNAP-25 knockout (KO) sampled tomogram subsets are shown in Figures 4 and 5, respectively. The results for all 101 tomograms are shown in Supplementary Figures 4 and 5. We analyzed the closest distance separating synaptic vesicles from the active zone in manually and automatically segmented Munc13- and SNAP-25-deficient synapses (Figure 4G-J; Figure 5G-J). Cumulative plots demonstrate a near overlap of measured vesicle distances, with inflection points typically deviating by less than ca. 5 nm between manual and automatic reconstructions (Figure 4G and H; Figure 5G and H). Next, we tested the sensitivity of SynapseNet to reveal synaptic vesicle docking deficits in mutant synapses by plotting the relative distribution of active-zone proximal vesicles normalized to active zone area (Figure 4I and J; Figure 5I and J). Munc13-1/2 DKO and SNAP-25 KO synapses exhibited a statistically significant reduction in vesicle occupancy within the 0-5 nm bin (Figure 4I: p=0.016; Figure fI: p=0.032), however, this reduction failed to reach statistical significance for the corresponding analysis of automatically segmented synapses (Figure 4J: p=0.127; Figure 5J: p=0.421). For both Munc13 and SNAP-25 mutants analyzed, the accumulation of non-docked vesicles within 5-20 nm of the active zone membrane in the smaller sampled subset of tomograms deviated from previously published data (Imig et al., 2014). Although manual reconstruction of Munc13-1/2 DKO and SNAP-25 KO synapses revealed a tendency towards increased vesicle occupancy of the 5-10 nm bin (Figure 4I and Figure 5I), this trend was exclusively detected within the 10-20 nm bin in the analyses performed with automatic segmentation results (Figure 4J and Figure 5J). Such discrepancies likely reflect the fact that only a smaller randomly selected subpopulation of synapses were sampled from the full datasets previously analyzed (Imig et al., 2014). Additionally, discrepancies between automatic and manual segmentations of synaptic vesicles or active zone that manifest within the range of one or two voxels (voxel size = 1.554 nm) can assign vesicles to neighboring bins. Despite the high stringency of this application, measurements derived from SynapseNet reliably reported the comparatively less severe docking phenotype exhibited by SNAP-25 compared to Munc13-1/2 -deficient synapses (Imig et al., 2014). Analogously, a comparative analysis of synaptic vesicle size demonstrated that automatic vesicle segmentation was also sensitive to increased vesicle diameter and volume upon Munc13-1/2 (Figure 4K-N) and SNAP-25 (Figure 5K-N) deletion. For Munc13-1/2 DKO synapses, manually and automatically segmented vesicles increased in diameter to 109.1% (Figure 4L) and 108.3% (Figure 4M) of controls, respectively. For SNAP-25 KO synapses, manually and automatically segmented vesicles increased in diameter to 115.1% (Figure 5L) and 114.5% (Figure 5M) of controls, respectively. For both Munc13- (Figure 4N) and SNAP-25-deficient (Figure 5N) synapses, these diameter changes translated to statistically significant increases in vesicular lumenal volume. Consistent with past studies (Imig et al., 2014), the extent of volume increase was correctly assessed by automatic segmentation to be greater for that of synapses lacking SNAP-25 (160.8% of controls) than for those lacking Munc13 (134.6% of controls).

**Figure 4:**
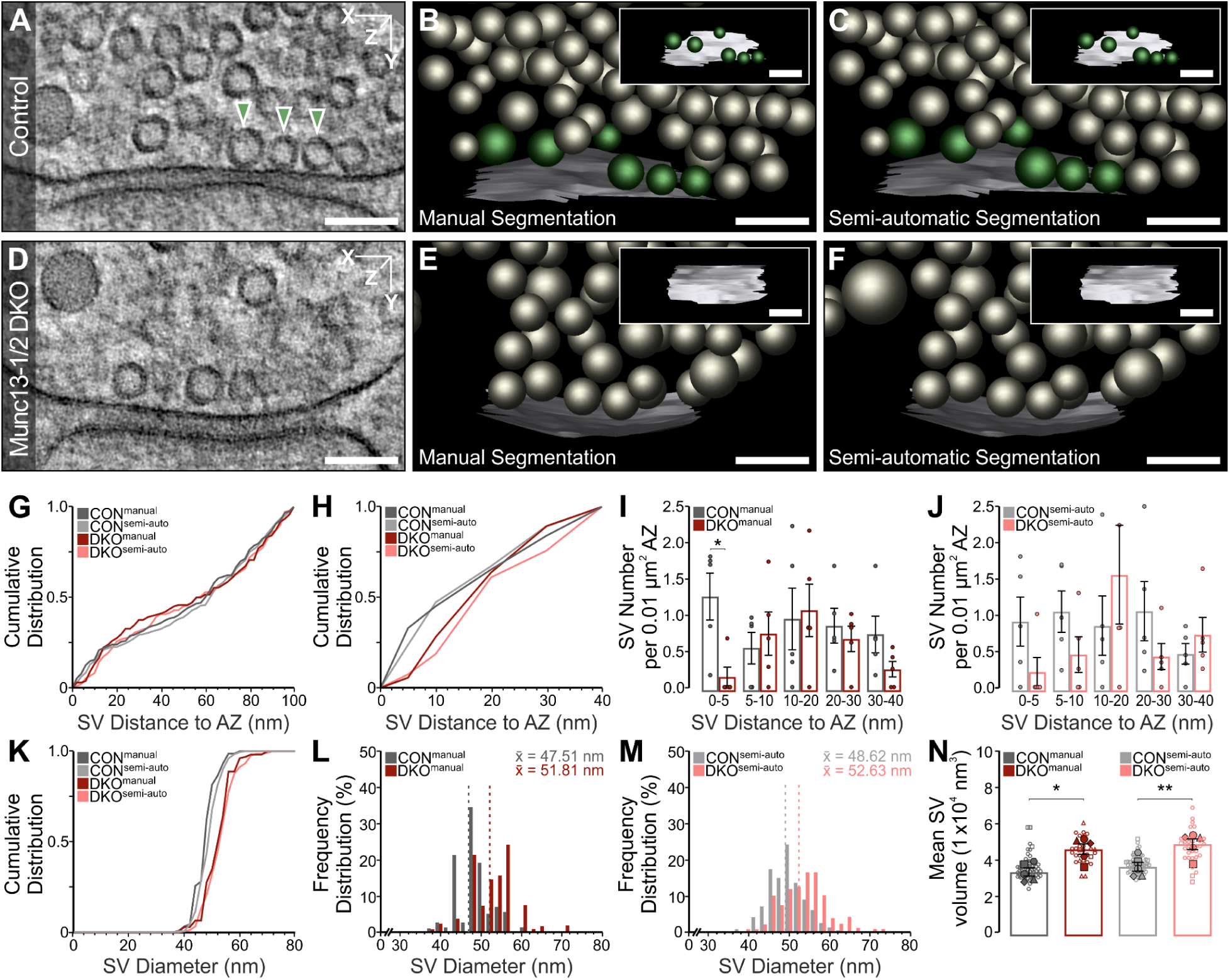
3D Electron tomographic analysis of synaptic vesicles in manually and automatically annotated Munc13-½ double knock-out (DKO) hippocampal Schaffer collateral neurons. **A, D.** Electron tomographic (ET) subvolume of control (CON) and Munc13-1/2 DKO (DKO) synapses. Docked synaptic vesicles (SVs) making active zone (AZ) contact within the displayed subvolume (arrowheads). **B, C, E, F.** 3D rendering of manual and automatic annotated synaptic profiles including orthogonal views of the AZ [AZ, white; docked SVs, green; non attached SVs, gray]. Analyses are based on manual and automatic annotation of SVs, as well as manual and semi-automatic segmentation of the AZ. **G, H.** Cumulative spatial distribution of SVs within 100 nm and 40 nm of the AZ. **I, J.** Mean SV number within 0–5, 5–10, 10–20, 20–30, 30–40 nm of the AZ normalized to AZ area. Data points indicate the SV number normalized to the AZ area of single ET subvolumes. **K.** Cumulative distribution of SV diameters within 100 nm of the AZ. **L, M.** Frequency distribution of SV diameters within 100 nm of the AZ. Dotted lines indicate the mean SV diameter (x̅). **N.** Scatterplot of the mean volume of SVs within 100 nm of the AZ. Filled data points indicate the mean SV volume of single ET subvolumes and empty data points with the same symbol shape indicate the volume of single SVs within the same ET subvolume. Values indicate mean ± SEM; ****p < 0.0001; ***p < 0.001; **p < 0.01; *p < 0.05. 5 ET subvolumes of CON and 5 of DKO synapses were analysed. The scale bars represent 100 nm.

**Figure 5:**
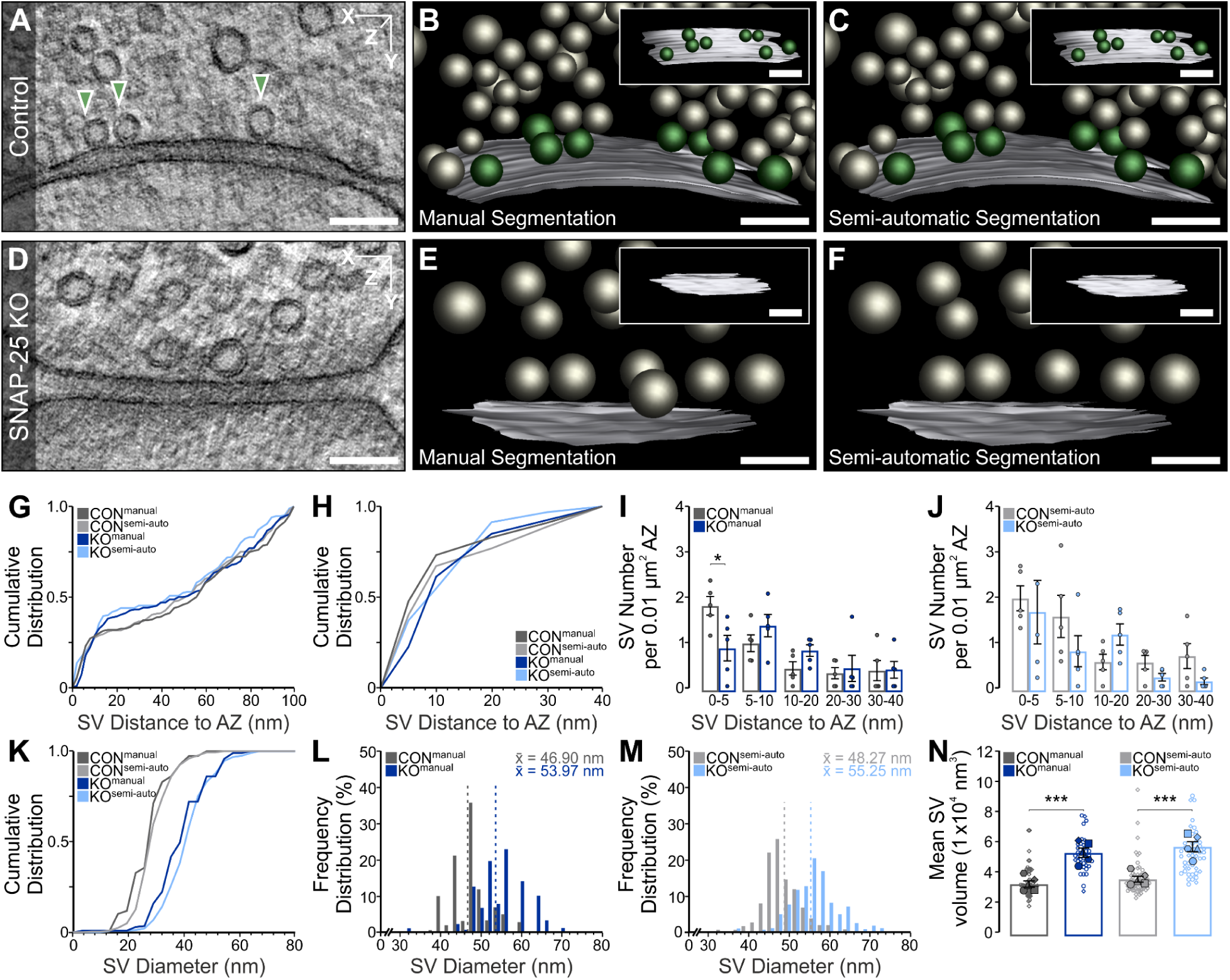
3D Electron tomographic analysis of synaptic vesicles in manual and automatic annotated SNAP-25 KO hippocampal Schaffer collateral neurons. **A, D.** Electron tomographic (ET) subvolume of control (CON) and SNAP-25 knock-out (KO) synapses. Docked synaptic vesicles (SVs) making active zone (AZ) contact within or outside of the displayed subvolume (arrowheads). **B, C, E, F.** 3D rendering of manual and automatic annotated synaptic profiles including orthogonal views of the AZ [AZ, white; docked SVs, green; non attached SVs, gray]. Analyses are based on manual and automatic annotations of SVs, as well as manual and semi-automatic segmentation of the AZ. **G, H.** Cumulative spatial distribution of SVs within 100 nm and 40 nm of the AZ. **I, J.** Mean SV number within 0–5, 5–10, 10–20, 20–30, 30–40 nm of the AZ normalized to AZ area. Data points indicate the SV number normalized to AZ area of single ET subvolumes (p 0-5 = 0.055556). **K.** Cumulative distribution of SV diameters within 100 nm of the AZ. **L, M.** Frequency distribution of SV diameters within 100 nm of the AZ. Dotted lines indicate the mean SV diameter (x̅). **N.** Scatterplot of the mean volume of SVs within 100 nm of the AZ. Filled data points indicate the mean SV volume of single ET subvolumes and empty data points with the same symbol shape indicate the volume of single SVs within the same ET subvolume. Values indicate mean ± SEM; ****p < 0.0001; ***p < 0.001; **p < 0.01; *p < 0.05. 5 ET subvolumes of CON and 5 of DKO synapses were analysed. The scale bars represent 100 nm.

We also used SynapseNet to perform an analysis of the active zones of inner ear ribbon synapses and compared it to the same analysis derived from manual annotations. Ribbon synapses occur in auditory and vestibular hair cells of the inner ear as well as in photoreceptors and bipolar cells of the retina. The active zones of these synapses contain the ribbon, an electron dense structure that tethers a halo of synaptic vesicles near the active zone membrane. Functions attributed to the synaptic ribbons include establishing a large pool of readily releasable vesicles and additionally aiding fast vesicle replenishment^49^. To demonstrate that SynapseNet can be used for rapid yet faithful analysis of ribbon synapses, we automatically segmented synaptic vesicles, ribbons, presynaptic densities, and active zone membranes in 88 tomograms with SynapseNet. In keeping with previous studies^11,38^, we defined three different vesicle pools: Ribbon-associated synaptic vesicles that are within a distance of 80 nm from the ribbon, membrane-proximal vesicles that are within a distance of 100 nm from the presynaptic density and 50 nm from the membrane, and docked vesicles that are within 100 nm from the presynaptic density and within 2 nm from the active zone membrane. We manually corrected the segmentation results using napari; particularly presynaptic densities had to be corrected. They were frequently misidentified by the model, likely due to their diffuse appearance. We then proof-read vesicle pool assignments in a second round of corrections, again using napari. Additionally, we annotated these structures in 33 tomograms manually with IMOD. To analyze the synaptic morphology, we determined the number of vesicles per pool and tomogram, the distances of ribbon-associated vesicles to the ribbon, the distances of membrane-proximal and docked vesicles to the presynaptic density and to the active zone membrane, and the synaptic vesicle diameters. The corresponding results are shown in Figure 6, which reports it for the set of tomograms with manual annotations, for the two analysis approaches, manual annotation and automatic segmentation followed by proof-reading. Both approaches yield very similar measurements for all reported quantities. A small discrepancy exists for the number of ribbon-associated synaptic vesicles per tomogram (Figure 6D), where manual assignment had about two more such vesicles per tomogram. This is most likely explained by the fact that the manual assignment was not based on a precise distance measurement, but only a rough estimate of the distance between vesicle and ribbon, leading to an inclusion of more vesicles than a stringent criterion based on a maximal distance. There is a difference in the cumulative distances in Figure 6F, but only for the different sets of tomograms (all vs. subset with manual annotation), most likely explained by different proportions of phenotypes in these sets, thus suggesting that SynapseNet will be able to distinguish those phenotypes. Another difference can be observed for the vesicle diameters (Figure 6F,G,J,K). Here, SynapseNet shows an approximately normal distribution, whereas the manual measurement has multiple distinct peaks. The likely explanation for this are artifacts from manual measurement (bias to select a radius from a discrete set for annotation). Alternatively, this could be explained by subsets of physiologically different vesicles, e.g due to different neurotransmitter filling state. In addition to the measurements here, Supplementary Figure 6 shows the analysis results for all tomograms, for two different stages of proof-reading (only correcting structure segmentation and also correcting vesicle pool segmentations and assignments). Here, we also found very similar distributions, showing that the automatic vesicle segmentation and assignment were already quite reliable. Overall, this analysis demonstrates that SynapseNet can be used to automate the very time consuming vesicle segmentation task for inner ear ribbon synapses, while providing good initial results for structure segmentations that can be corrected, replacing the need for fully manual annotation. It will thus enable the analysis of different phenotypes in inner ear ribbon synapses in the future.

**Figure 6:**
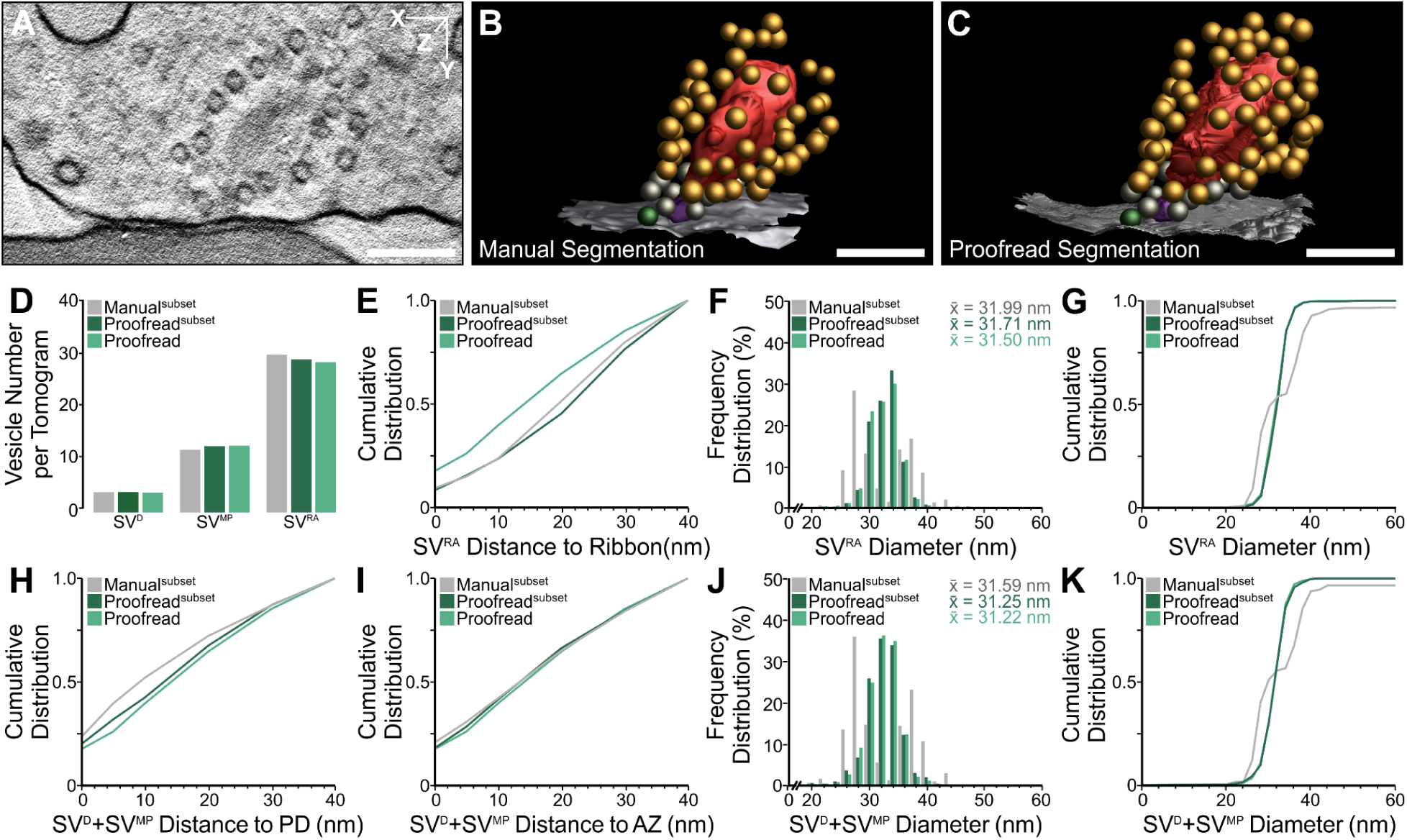
Analysis of ribbon synapses of cochlear inner hair cells in the inner ear. **A-C.** Electron tomographic subvolume and 3D rendering of synaptic vesicle (SV) pools for manual and semi-automatic segmentation [active zone (AZ), white; docked SVs, green; membrane-proximal SVs, gray; ribbon-associated SVs, orange; ribbon, red; presynaptic density (PD), magenta]. **D-K.** A subset of manual and proofread segmentations (n=33), and a larger dataset of proofread segmentations (n=88) were analysed. **D.** Average number of docked SVs (SV^D^), membrane-proximal SVs (SV^MP^), and ribbon-associated SVs (SV^RA^) per tomogram. **E.** Cumulative spatial distribution of SV^RA^ within 40 nm of the ribbon. **F,G.** Frequency distribution of SV^RA^ diameters with the mean SV diameter (x̅) and cumulative distribution of SV diameters. **H,I.** Cumulative spatial distribution of SV^D^+SV^MP^ within 40 nm of the PD and the AZ. **J,K.** Frequency distribution of SV^D^+SV^MP^ diameters with the mean SV diameter (x̅) and cumulative distribution of SV diameters. Values indicate the mean diameters. The scale bars represent 200 nm.

### SynapseNet Tool

We built a graphical user interface to make SynapseNet’s segmentation and analysis functionality available to researchers without programming expertise. It is implemented as a napari^37^ plugin that provides widgets for:

- Automatic segmentation of synaptic structures. This widget provides synaptic vesicle segmentation with three different models (for 2D electron micrographs, room-temperature electron tomography, and cryogenic electron tomography), active zone segmentation, ribbon structure segmentation, mitochondrion segmentation, and synaptic compartment segmentation.
- Distance measurements. This widget can measure distances between vesicles and another segmented object, for example the active zone, as well as measure pairwise distances between vesicles. The distance measurements can be displayed within napari and can be exported to a table.
- Morphological measurements. This widget enables the measurement of vesicle morphology (radii and intensity statistics) and object morphology (surface area and volume).
- Synaptic vesicle pool assignments. This widget enables assigning vesicles to pools based on the distances and/or morphology measurements and user-defined criteria.

The combined functionality of our plugin enables the extraction of data for analysis in a fast manner requiring minimal intervention. We implemented SynapseNet’s functionality, including the plugin, in a well-documented python library that computationally experienced users can use to automatically process large datasets. This also includes code and examples for training a segmentation network for a new task, given a dataset with annotations, and for adapting one of the pre-trained networks to a new imaging modality or otherwise different condition (without annotations) using domain adaptation. We further provide functionality to export segmentation results to the IMOD data format for analysis in 3DMOD.

## Discussion

We introduced SynapseNet, a tool for automatic synapse reconstruction in electron microscopy. It provides a large annotated dataset and methods for automatic segmentation of vesicles and other synaptic structures, and a domain adaptation method to improve segmentations for different imaging modalities. We demonstrated its utility in two example analyses and implemented a python library and graphical user interface to apply it in research applications.

This contribution marks the first comprehensive approach for automatic synapse reconstruction from electron micrographs. We foresee that it will significantly speed up the analysis of synaptic ultrastructure, enabling neuroscientists to study much larger data samples and derive novel insight into the morphological organization of synapses.

In the segmentation evaluation we focused on synaptic vesicles, testing different electron microscopy modalities and demonstrating good segmentation results for a broad range of settings, including volumetric and 2D data, especially when combined with domain adaptation. We also found some limitations, where the models did not work for data that is too different from the training data, particularly for those with low contrast. We evaluated the models for other segmentation tasks – active zones, mitochondria, ribbon synapse structures, and synaptic compartments – less extensively. They provided good results for the conditions we tested them in and can potentially also be adapted to different modalities through domain adaptation.

SynapseNet is a big step towards the automatic analysis of synapses based on electron microscopy. However, in the current state it does not enable full automation for any such analysis task. Depending on the data, segmentation results may be incomplete, even after domain adaptation, with, for example,missing synaptic vesicles or active zones, or false splits in synaptic compartments. Nevertheless, it can still speed up analysis massively in these cases by enabling semi-automatic segmentation. Our user-interface, which inherits the proof-reading capabilities of napari, enables segmentation correction through painting as well as merging and erasing of objects. Segmentation results can also be exported to IMOD, which has its own correction functionalities. The python library makes SynapseNet compatible with most other tools for segmentation correction or analysis. In cases where our models do not work for a given segmentation task, or where we do not provide a model for the task, our training functionality can be used if provided sufficient annotated data.

To automate synaptic segmentation tasks further, we plan to collect more training data, including for 2D images from serial section transmission electron microscopy and for other synaptic structures such as actin (in cryo electron tomography), dense core vesicles, endo- and exosomes, microtubules, or smooth endoplasmic reticulum. Currently, SynapseNet focuses on the presynapse, and we plan to also extend it to the postsynapse. The organelles in the postsynapse are not as well characterized as in the presynapse yet, but studies based on electron microscopy^14^ and super-resolution microscopy^50^ have shown the connection between postsynaptic morphology and function. Extension to the postsynapse would thus enable a more comprehensive analysis of its morphology, including analysis of the postsynaptic density, which is visible in electron micrographs given appropriate sample preparation. Further, we plan to develop a shared architecture for the different segmentation tasks that could improve segmentation performance by exploiting similarities and spatial context between different structures. Such a model could also enable better generalization to further improve domain adaptation. This development could profit from recent work on vision foundation models^7,51^. It would further require learning a multiscale data representation to segment structures of different sizes, e.g. synaptic vesicles and compartments, with a shared model. Fortunately, such multiscale models have been studied in other contexts^52^ and could potentially be extended by us. Another important goal is the integration of our tool with methods for protein identification in cryo electron tomography^53,54^ to enable the joint analysis of morphology and molecular information in the synapse.

## Materials and Methods

### Sample Preparation and Data Acquisition

In this section we list all datasets that were used for model training, evaluation, for domain adaptation, and for example analysis. We used data from different electron microscopy modalities and different sample preparation techniques. Most of the data is room temperature electron tomography, but we also included cryogenic electron tomography and serial section transmission electron microscopy. The data covers different synapse types from different organisms. It includes both previously published and yet unpublished data. We standardized data and annotation formats, see <u>Data Annotation</u> for details. Supplementary Table 1 shows a concise overview; for each data subset we give the abbreviation used for it throughout the manuscript in the beginning.

### Single-Axis TEM Tomo & Chemical Fixation

We used 152 room-temperature single-axis TEM tomograms from Schaffer collateral and mossy fiber synapses in organotypic hippocampal slices for supervised training of synaptic vesicles, mitochondria and active zones, evaluation of synaptic vesicle and active zone segmentation, and for validation of the approach to detect changes in presynaptic morphology upon deletion of presynaptic proteins (Figures 4 and 5). The tomograms were generated in two published studies^16,17^. Organotypic slices were prepared from the hippocampi of neonatal mice according to the interface protocol^55^ and vitrified after 28 days *in vitro* in culture medium supplemented with 20% (w/v) bovine serum albumin using an HPM100 (Leica) high-pressure freezing device. We used 23 tomograms resulting from chemically-fixed material, which were also published in (Maus et al., 2020)^16^. For these tomograms, wild-type animals at postnatal day 28 were transcardially perfused under deep anesthesia, first with 0.9% sodium chloride solution, and then one of two fixatives (Fixative 1: Ice-cold 4% paraformaldehyde, 2.5% glutaraldehyde in 0.1 M phosphate buffer^15^; Fixative 2: 37° C 2% paraformaldehyde, 2.5% glutaraldehyde, 2 mM CaCl_2_, in 0.1 M cacodylate buffer^56^). Brains were rinsed and sectioned coronally through the dorsal hippocampus in an ice-cold 0.1 M phosphate buffer using a VT 1200S vibratome (Leica) (step size 100 µm; amplitude 1.5 mm, speed 0.1 mm/sec). Hippocampal CA3 subregions were excised using a 1.5 mm diameter biopsy punch and high-pressure frozen on the same day in 20% (w/v) bovine serum albumin using an HPM100 (Leica) high-pressure freezing device. For both sample preparations, automated freeze-substitution was performed as previously described in (Imig and Cooper, 2017)^57^. Tomograms were collected using a 200 kV JEM-2100 (JEOL) transmission electron microscope equipped with an 11 MP Orius SC1000 CCD camera (Gatan). Tilt-series (tilt range +/-60°; 1° angular increments) were acquired at 30 000x magnification using SerialEM^58^. Tomographic reconstructions were generated using weighted back-projection with etomo^25^.

### Dual-Axis TEM Tomo

We used 114 unpublished room-temperature dual-axis TEM tomograms generated from primary hippocampal neuron cultures and from organotypic hippocampal slices for supervised training and evaluation of synaptic vesicle segmentation. Neuronal monolayer cultures from newborn C57/BL6J mice were cultured on astrocyte-bearing sapphire discs as previously described by (Watanabe et al., 2014)^59^. Organotypic slices were prepared from the hippocampi of neonatal mice according to a modified roller-tube slice culture protocol (Imig et al., 2020)^12^. Primary neuron cultures and organotypic slices were vitrified at 14 and 28 days *in vitro*, respectively, without added cryoprotectants using an EM ICE (Leica) high-pressure freezing device. Tomograms were collected using a 200 kV Talos F200C G2 scanning/transmission electron microscope equipped with a Ceta CMOS camera (Thermo Scientific). Orthogonal tilt series (tilt range +/-60°; angular increment 1°) were acquired at either 57000 x magnification (neuron culture) or 36000 x (organotypic slice culture) using SerialEM^58^. Tomographic reconstructions were generated using weighted back-projection with etomo^25^.

### STEM Tomo

We used 48 unpublished room-temperature dual-axis STEM tomograms generated from hippocampal Schaffer collateral, mossy fiber, and perisomatic synapses for supervised training and evaluation of synaptic vesicle, active zone, mitochondria, and compartment segmentation. Tomograms were acquired using a 200 kV Talos F200 G2 scanning/transmission electron microscope (Thermo Fisher Scientific) equipped with a Model 2040 dual-axis high-angle tomography holder (Fischione). STEM imaging (format 2048 x 2048; pixel size 0.869 nm; dwell time 7 µs/pixel) was performed in microprobe mode with a beam semi-convergence angle of 3.6 mrad. Images were formed with an on-axis brightfield detector using a collection angle of 10 mrad. Continuous tilt series were acquired (tilt range +/-60°; angular increment 1°) from orthogonal axes in low-dose and dynamic focus modes using SerialEM^58^ and reconstructed with weighted back-projection implemented by etomo^25^.

### IER

We used room temperature electron tomograms of murine and rat inner ear ribbon synapses for training and evaluation of ribbon synapse structures. We used 6 tomograms from mice from two studies^11,38^, as well as 3 additional tomograms that were acquired following the same protocol as (Chakrabarti et al., 2022)^11^ but that were not included in the publication. Further, we used 19 tomograms from rats from (Michanski, Kapoor et al., 2023)^39^. We included additional 88 newly acquired tomograms from mice, for which the organs of Corti were acutely isolated, high-pressure frozen with a HPM100 and freeze substituted as described in (Chakrabarti et al., 2018)^38^. For these tomograms, single-axis tilts were acquired with a 200 kV JEM-2100 (JEOL) transmission electron microscope equipped with a 11 MP Orius SC 1000 CCD camera (GATAN) and a JEM 2100 Plus (JEOL) equipped with a 20 MP XAROSA bottom-mount CMOS TEM camera (EMSIS), mainly tilting from – to +60° using the Serial-EM software package^58^. Acquired tilt-series were generated into tomograms utilizing etomo from the IMOD software package^25^. These tomograms were also used for the domain adaptation evaluation and for the analysis of vesicle pools in the inner ear ribbon synapses in Figure 6.

### EH

We used 44 room temperature electron tomograms of murine endbulb of Held synapses to evaluate vesicle segmentation domain adaptation. For these tomograms, Parasagittal vibratome sections of the anteroventral cochlear nucleus were freshly prepared and instantly high-pressure frozen with a HPM100 and subsequently freeze substituted as described in (Hintze et al., 2021)^60^. The tomograms were acquired as single-axis tilts at a JEOL JEM 2100 Plus equipped with an 20 MP XAROSA bottom-mount CMOS TEM camera (EMSIS) from – to + 60° using the Serial-EM software package^58^. Acquired tilt-series were generated into tomograms utilizing etomo from the IMOD software package^25^.

### Cryo

We used 22 cryogenic electron tomograms to evaluate vesicle domain adaptation. For these tomograms, primary rat hippocampal neurons were cultured on EM grids (R2/2 SiO2 film on Au mesh, Quantifoil) coated with poly-L-lysine. Prior to plunge-freezing the cells (DIV 14-18) were incubated for approximately 5 minutes in a Tyrode solution containing 5% (v/v) glycerol acting as a cryoprotectant. Electron transparent lamellae were produced using focused ion beam (FIB) milling (Aquilos 2 Cryo-FIB, ThermoFischer Scientific). Tomograms were collected with a 300 kV Krios G4 Cryo TEM microscope (ThermoFischer Scientific) on regions of interest exhibiting typical synaptic markers (e.g. presence of synaptic vesicles) previously identified on low magnification transmission electron microscope (TEM) overviews. Tomograms were reconstructed using a weighted-back projection algorithm implemented in the IMOD software^25^. Tomograms were then deconvolved using a Wiener-like filter implemented in the TOM toolbox^61^.

### Frog

We used 402 transmission electron micrographs of synapses of a frog to evaluate 2D vesicle domain adaptation. For this data frog (*Rana pipiens*) cutaneous pectoris muscles were dissected and mounted in sylgard-lined chambers, in a frog Ringer buffer (115 mM NaCl, 2 mM KCl, 1.8 mM CaCl_2_, 2.4 mM NaHCO_3_, pH 7.2). Samples were fixed with 2% glutaraldehyde in frog Ringer, at -2°C for 40-60 minutes, followed by washing, post-fixing with 2% OsO_4_ in PBS, dehydration through a series of ethanol solutions and propylene oxide, and embedding in Epon resin. A fraction of the vesicles was labeled with precipitated di-amino-benzidine, using a photo-oxidation procedure. The blocks were then sectioned (80-90 nm thickness), and were post-stained with 2% uranyl acetate in a 1:1 ethanol-water mixture (1 minute incubation). Images were acquired using a CM-10 Philips electron microscope, using film negatives. These were enlarged 2.69-fold by transferring to photographic paper (Eastman Kodak Company, Rochester, NY). The paper was scanned at 300 dpi, to obtain the final images. The data were published in (Rizzoli and Betz)^43^ and were obtained more than 20 years ago.

### 2D TEM

We used 13 transmission electron micrographs (large 2D images, ca. 5,000 x 5,000 pixels) of Schaffer collateral and mossy fiber synapses in organotypic hippocampal slices for vesicle domain adaptation. The tomograms were published as part of the study^16^. Organotypic slices were prepared from the hippocampi of neonatal mice using the interface protocol (Stoppini et al., 1991; Imig et al., 2014)^17,55^ and vitrified after 28 day *in vitro* in culture medium supplemented with 20% (w/v) bovine serum albumin using an HPM100 (Leica) high-pressure freezing device. Automated freeze-substitution and epoxy embedding was performed as previously described (Imig and Cooper, 2017)^57^. Ultrastructural analysis was performed on 60 nm-thick sections postcontrasted with 1% aqueous uranyl acetate and Reynold’s lead citrate. Electron micrographs were acquired at 20 000 x magnification (pixel size = 0.592 nm) with an 80 kV LEO 912-Omega transmission electron microscope (Zeiss) equipped with a slow scan dual-speed CCD camera “Sharpeye” (Tröndle, Moorenweis, Germany).

### Munc13 / SNAP-25

We used 101 tomograms for the analysis of Munc13 / SNAP-25 knockouts (Figures 4 and 5). These tomograms were acquired by (Imig et al. 2014)^17^. The sample preparation and data acquisition followed the same protocol as for the **Single-Axis TEM Tomo** data (see above).

## Data Annotation

### Synaptic vesicles

We made use of 337 tomograms (datasets: Chemical fixation, Single-Axis TEM Tomo, Dual-Axis TEM Tomo, STEM Tomo; see previous section) with synaptic vesicle annotations for training and evaluation. For these tomograms, initial vesicle annotations were exported from IMOD, either available from the respective study or newly annotated for unpublished data. For unpublished data the IMOD annotations were partially created based on segmentations from an initial version of our vesicle model (see also below), which were then exported to IMOD and proofread. In most tomograms the synaptic vesicles were only partially annotated, often only in proximity of the active zone(s). Furthermore, they were annotated as spheres, which only approximate the true vesicle shape. Our supervised training procedure requires dense annotations, i.e. all vesicles in a (cropped) tomogram have to be annotated. Annotations should also adhere to the exact position of vesicular membranes to avoid the introduction of shape biases. We post-processed the IMOD annotations to resolve these issues. First, we produced the crops tightly enclosing the IMOD annotations for all tomograms and trained an initial model on this dataset. We then selected a subset of tomograms, for which the crops contained almost dense annotations and for which the (spherical) annotations had a good fit to the actual vesicle membranes. We reran this network on all tomograms, and corrected the vesicle annotations via a seeded watershed, using the model predictions as heightmap. We then included new tomograms, for which the shapes now matched better, still restricted to almost dense annotations, into an extended training set. We trained a new version of the network on this, repeating this process three times in total. To obtain the final version of the training set, we then ran prediction on all tomogram crops, including those with sparse annotations, and added vesicles from the network predictions that did not intersect with a vesicle annotation to the ground-truth. We visually inspected the result and found almost no errors. Our final training data for vesicle segmentation contained 117,112 annotations, 80,225 of these derived from manual annotations and the remainder added based on automatically segmented vesicles. We split this data into 269 tomograms with 66,292 annotated vesicle for training and 68 tomograms with 51,420 vesicles for testing.

### Active zones

We exported the IMOD annotations for 209 active zones from 183 tomograms (datasets: Single-Axis TEM Tomo, Chemical Fixation, and STEM Tomo), which were either prepared for previous studies or newly annotated. The annotations contained some artifacts caused by the IMOD export. We found that training a network to segment active zones was still feasible based on them, but further post-processing would likely improve results. We manually proofread the tomograms used for evaluating the active zone segmentation (Supplementary Figure 2B) to remove these artifacts. We split this data into 137 tomograms with 161 active zones for training and 46 tomograms with 48 active zones for testing.

### Mitochondria

We exported the IMOD annotations for 90 mitochondria from 29 tomograms. We had to manually curate these annotations in order to remove artifacts resulting from the export. We did not use a test split due to the small number of tomograms and mitochondria.

### Ribbon synapse structures

We exported the IMOD annotations for the synaptic ribbon, the presynaptic density, and the active zone membrane for an initial set of 49 tomograms, composed of the tomograms from previous studies and 21 of the newly acquired tomograms. Based on these tomograms we trained an initial segmentation model and then reran prediction for the other 67 tomograms from the newly acquired dataset. For these, the predictions were manually corrected and then included as additional training data. Note, that we did not evaluate the segmentation of the active zone membrane, as these were not performed following a strict procedure, making it difficult to consistently evaluate predictions. The corrected active zone segmentations were used for the analysis in Figure 6.

### Synaptic compartments

We annotated 289 synaptic compartments in 28 STEM tomograms using μSAM^7^, a tool for interactive and automatic microscopy segmentation based on Segment Anything^51^. We first annotated compartments in 2D on virtual tomogram sections, using the default ViT-B Segment Anything Model in μSAM. We then finetuned this model on these annotations, significantly improving its segmentation performance for the task, and used this model to interactively annotate the compartments in 3D. We did not use a test split due to the small number of tomograms and compartments.

### Vesicle segmentation (domain adaptation)

We did not require any annotations to perform the domain adaptation training for synaptic vesicles. However, to quantitatively evaluate these results we needed annotations for a few tomograms. These were obtained for the respective datasets as follows:

- IER: We corrected the predictions of an initial domain adapted model in crops of 12 of the newly acquired tomograms. The corrected crops contained 1,706 annotated vesicles. We excluded these tomograms from the training data for domain adaptation.
- EH: We annotated all vesicles in a presynaptic compartment for 5 tomograms in IMOD, also providing an annotation for the respective compartment. This resulted in 949 annotated vesicles. The respective tomograms were excluded for domain adaptation training and the segmentation evaluation was restricted to the compartment masks.
- Cryo: We obtained 2 tomograms with binary annotations for synaptic vesicle membranes from MemBrain^34^ that were then corrected with the Amira software (Thermo Fisher Scientific). We then created an instance segmentation via a seeded watershed, using centroids of the connected components of the annotations as seeds and their distance transform as heightmap. The result was manually corrected in napari.
- 2D TEM: We used annotations for four tomograms from IMOD that were produced for previous analysis of this data. These annotations were concentrated around active zones, and we manually delineated masks around the annotations that were used to restrict the segmentation evaluations.
- Frog: We did not quantitatively evaluate this data and thus did not make use of any vesicle annotations.

#### Other Tools for Synaptic Reconstruction

Here, we briefly describe the tools offering segmentation functionality for synapses in electron micrographs known to us. Out of these tools we only found CryoVesNet^30^ that targets one of the segmentation tasks we address, that is easy-to-use, and that scales to processing large datasets. Thus, we use it for benchmarking in our experiments.

Multiple tools address synaptic vesicle segmentation in cryogenic electron tomography. These include tools using template matching^23,29^, which rely on parameter tuning or other manual intervention, as well as a high signal-to-noise ratio. More recently, deep learning based methods have been introduced for this task. This includes CryoVesNet^30^, which provides a 3D segmentation network trained based on tomograms with vesicle annotations, and VesiclePicker^31^, which uses the Segment Anything Model^51^ to automatically segment synapses. The latter is tightly integrated with cryoSPARC^62^, which is a popular software for cryogenic electron microscopy, although not widely used in other fields. It thus cannot be easily used for synaptic vesicle analysis in a variety of settings. The tools TomoSegMemTV^35^ and MemBrain^34^ provide segmentation of plasma membranes. This makes them useful for picking membrane-bound proteins, however, analyzing the morphology of individual objects would require additional steps to identify these objects. Dragonfly^36^ is popular for segmentation tasks in cryogenic electron tomography, but lacks pretrained models for specific tasks, resulting in a need for extensive user annotations for training.

Synaptic vesicle segmentation and similar tasks have also been addressed for room-temperature electron microscopy. Imbrosci et al.^33^ implement vesicle detection in 2D. This solution does not enable morphology analysis or distance measurements because it only provides counts and centroid positions. VeSElecT^27^ implements vesicle segmentation in 3D based on classical image analysis. It is implemented as a Fiji plugin that requires manual parameter tuning, and can thus not be applied automatically to large datasets. To our knowledge, there are no methods for segmenting active zones, synaptic compartments, or ribbon synapse structures. MitoNet^8^ provides a general-purpose solution for mitochondria segmentation in electron microscopy. However, we did not compare to it due to the limited number of mitochondrion annotations in our dataset.

#### Supervised segmentation

We implemented segmentation functionality for synaptic vesicles, active zones, mitochondria, ribbon synapse structures, and synaptic compartments. We trained 2D and 3D segmentation networks for vesicles and 3D segmentation networks for the other structures, using the 2D^3^ and 3D UNet^4^ implementations of torch-em^63^, which is based on PyTorch^64^. Unless specified otherwise, we follow the same approach for all segmentation tasks: The networks predict foreground and boundary probabilities for the respective structure. They were trained with the respective annotated data (see <u>Data Annotation</u> for details). The datasets were divided into a training, validation, and test split. We used the negative Dice coefficient as the loss function. After prediction, structure-specific post-processing is applied to the network outputs to obtain the segmentation result, see below for details. We used the following training hyperparameters:

- Optimizer: AdamW^65^ with an initial learning rate of 10^-4^ and PyTorch defaults otherwise.
- Learning Rate Decay: The learning rate was decreased by a factor of 0.5 when the validation loss plateaued for 5 epochs.
- Iterations: The networks were for 100,000 iterations and the checkpoint of the epoch with the lowest validation loss was exported.
- Network architecture: A UNet with 4 levels, 32 initial features and an increase of the number of features by a factor of 2 in each level. The feature representations are downsampled spatially by a factor of 2 after each level via max pooling in the encoder and upsampled by the corresponding factor in the decoder via interpolation. In the case of the 3D UNets, the representations are downsampled only in the image plane after the first level, followed by isotropic sampling afterwards. This approach was chosen to account for the smaller image dimension across the depth axis and the missing wedge effect.
- Patch shapes:
  - Synaptic vesicles (2D): 256 x 256 pixels
  - Synaptic vesicles (3D): 48 x 256 x 256 voxels
  - Active zones: 48 x 256 x 256 voxels
  - Mitochondria: 32 x 256 x 256 voxels
  - Ribbon synapse structures: 64 x 512 x 512 voxels
  - Synaptic compartments: 64 x 384 x 384 voxels

Furthermore, the data was binned by a factor of 2 for mitochondria segmentation and by a factor of 4 for synaptic compartment segmentation. Consequently the data has to be binned by the same factor in prediction. We have stored the average (effective) pixel or voxel sizes for all training sets in our python library and by default resample the input data to the respective resolution to handle differences in the resolution of the training and user data.

We implemented the following post-processing procedures to obtain synaptic structure segmentations based on the respective network predictions:

- Synaptic vesicles: Analyzing individual vesicles requires an instance segmentation, which we obtain by applying a seeded watershed to the network’s foreground and boundary predictions. In more detail:
  - We compute the distances to the boundary predictions thresholded at a value of 0.5 with the euclidean distance transformation.
  - We threshold these distances at a value of 8 and compute the connected components to obtain the *seeds*.
  - We compute the distance to the nearest of these components, using the euclidean distance transform, to obtain the *heightmap*.
  - We threshold the foreground predictions at a value of 0.5 to obtain the *mask*.
  - We perform a seeded watershed using the *heightmap*, *seeds* and *mask* to obtain the vesicle segmentation.
  The approximate spherical shape enables recovering individual vesicles even with imperfect boundary predictions through this procedure, as seed components can be recovered by the distance based seed computation for “holes” in the boundary predictions.
- Active zones: The active zones are spatially well-separated, as such, it is not necessary to produce an instance segmentation, which can be trivially obtained by applying connected components later. We thus only threshold the network predictions at a threshold of 0.5, remove small connected objects with a size smaller than 500 pixels, and return the resulting binary segmentation.
- Mitochondria: To analyze individual mitochondria we obtain an instance segmentation based on the network’s foreground and boundary predictions. Here, we follow a similar approach compared to the vesicle segmentation, however, choose a lower value of 0.25 for the initial boundary threshold and a seed distance value of 6 pixels.
- Ribbon synapse structures: The ribbon and presynaptic density are spatially well-separated, hence, we return a binary segmentation. These segmentations are filtered by selecting the connected component with the most surrounding synaptic vesicles (derived from a vesicle segmentation) for the ribbon and selecting the component closest to the ribbon for the presynaptic density.
- Synaptic compartments: For synaptic compartments the network predicts a boundary channel and a channel that regresses the distance to the boundary that is used to compute an auxiliary loss. Based on the boundary predictions, we segment individual compartments by segmenting large compartments individually in 2D using a seeded watershed based on the distance to the boundary and then merging 2D compartments across the depth axis via a minimum cost multicut problem^66^ with costs derived from component overlaps.

We implemented the post-processing using the euclidean distance transform from SciPy^67^ and the connected component and watershed functions from scikit-image^68^. In addition, we used custom image analysis functionality from our library, <u>elf</u>.

#### Domain adaptation

The field of domain adaptation^69^ studies methods to make machine learning models more robust to so-called domain shifts of the input data between the training set (called *source domain*) and new test data (called *target domain*). Domain shifts are a common problem in biomedical images. For electron micrographs of synapses, domain shifts can arise due to different electron microscopy modalities, different sample preparation, and/or different synapse types in different animals, all resulting in a distribution shift of the image data. To address the domain adaptation problem, we implemented an algorithm based on the mean teacher method^40^. This method was initially introduced for semi-supervised classification and has previously been extended to domain adaptation for classification^41^ and to segmentation, including in our own prior work^42^. We address unsupervised domain adaptation, i.e. our method does not require annotations in the target domain.

An overview of our domain adaptation algorithm is shown in Supplementary Figure 7A. It works as follows:

- The starting point is a model (2D or 3D UNet) trained via supervised learning on the source domain, see also <u>Supervised segmentation</u>.
- The architecture of this model is duplicated to obtain a *teacher* and a *student* model. The weights of both models are initialized with the pretrained model’s weights.
- The student model is then trained via mini-batch stochastic gradient descent, using so-called pseudo labels, which are derived from the predictions of the teacher model. A single training iteration proceeds as follows:
  - A batch of images with the chosen patch shape is sampled from the target domain.
  - Augmentations sampled from a distribution of image transformations are applied to these images, keeping the original version of images.
  - The teacher model is applied to the augmented images. The gradients are not computed for this step.
  - A confidence mask *m_c_* is computed by selecting pixels in the teacher prediction that have a probability larger than the *confidence threshold t_c_*, or smaller than 1 - *t_c_*. The corresponding pixels are set to 1 in the mask, other pixels are set to 0. We used *t_c_* = 0.75 for all experiments.
  - The student model is applied to the original images.
  - The loss is computed between the predictions of the teacher, called pseudo-labels, and the predictions of the student. Only values that are positive in *m_c_* are taken into account for the loss computation. The negative dice score is used as the loss function.
  - The parameters of the student are updated using ADAM (same hyperparameters as before).
  - The weights of the teacher model *w_t_* are updated according to an exponential moving average of the student weights *w_s_*: *w_t_ =* α *w_t_ +* (1 - α) *w_s_*. We used α *=* 0.999 for all experiments.

We used Gaussian blur with a bandwidth sampled from the range [0, 2.5] and additive Gaussian noise with a noise scale (standard deviation) of [0%, 15%] (uniform sample) of the data range. Otherwise we used the same hyperparameters as in <u>Supervised segmentation</u>.

The motivation behind this approach is to only learn from the certain predictions of the teacher model by using the confidence threshold for loss masking. Due to the gradual update of the teacher weights through the exponential moving average, the teacher improves on the target domain, leading to improved pseudo labels and higher confidences, hence, also expanding the fraction of the data used for loss computation. Note that this approach can only be successful if the initial predictions of the teacher, corresponding to the predictions of the model trained on the source domain, find at least parts of the structures of interest. If they completely miss the structures of interest and/or find other structures with high confidence, then the domain adaptation will fail. Note that such failures can only be reliably detected by visually inspecting the segmentation results after adaptation. Such checks are important before deriving measurements for any data from segmentation results of a domain adapted network. The approach described here is an extension of our previous work on probabilistic domain adaptation^42^. The major differences are:

- We extended the approach to 3D segmentation.
- We used a regular UNet instead of a probabilistic UNet^70^, which was used for a more complex estimate of the confidence mask. Using a regular UNet simplified the overall approach, facilitated the extension to 3D, and did not result in noticeable differences in the domain adaptation outcomes according to preliminary experiments.
- We used a two-stage domain adaptation approach where a pretrained source model is adapted to the targeted domain. In our previous work we investigated different settings including joint training on source and target domain. We chose the two-stage approach here because it is simpler to apply in practice.

#### Segmentation metrics

We used standard metrics to evaluate the segmentation tasks. For synaptic vesicle segmentation we reported the F1-Score measured at the intersection over union of 50%. To compute this metric, the overlap between all objects in the segmentation and the corresponding annotations are measured. Object pairs with an overlap larger than 50% are counted as true positives (TP), objects in the segmentation that were not matched are counted as false positives (FP), and objects in the annotation that were not matched are counted as false negatives (FN). Therefore, the F1-Score is defined as: F1 = 2 * TP / (2 * TP + FP + FN). For binary segmentation tasks (ribbon synapse structures, active zone) we reported the Dice coefficient, which is a measure for the overlap between two binary sets, in our case the segmentation *s* and the annotation *a*. It is defined as 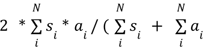). Here, *i* is the flattened pixel or voxel index, and *N* is the number of pixels / voxels. For multiple images or tomograms we reported the average evaluation score (F1 or Dice) and indicated the standard deviation when five or more images or tomograms were used for the corresponding evaluation experiments. For the evaluation of the active zone segmentation, we further post-processed both the annotations and predicted segmentations by intersecting them with a thresholded Sato ridge filter^71^ in order to ‘snap’ them to the presynaptic membrane. We then skeletonized the result to remove artifacts, and expanded the obtained line again via a morphological dilation and closing operation. This approach produced a more consistent overlap of predictions and annotations where the active zone was correctly predicted.

#### Synaptic analysis

For the MUNC13-1/2 DKO / SNAP25 KO tomograms (Analysis in Figure 4, 5) we first manually annotated 5 tomograms for each knock-out and the corresponding control (20 tomograms in total) using IMOD. Synaptic vesicles were annotated by exporting predictions from SynapseNet to IMOD and correcting them. The active zones were manually annotated. We then measured the shortest distance between the active zone and each synaptic vesicle using the mtk program, which is part of IMOD. In addition, vesicle diameters and active zone surface areas were extracted from segmented models using imodinfo. We further distinguished vesicles into non-attached and docked phenotypes; a vesicle was defined as docked when there was no measurable distance between the outer leaflet of the vesicle and the inner leaflet of the lipid bilayer (i.e. when the dark pixels corresponding to the vesicular outer leaflet were contiguous with those of the inner plasma membrane leaflet). Based on the voxel size of 1.554 nm, these docked vesicles fall into the 0-2 nm bin. The number of vesicles in discrete bins (i.e. 0-2 nm, 0-40 nm, and 0-100 nm from the active zone membrane) were normalized to the measured active zone surface area and reported as the number of vesicles per 0.01 µm^2^. We redid these steps automatically with SynapseNet for all 101 available tomograms. We first applied automatic synaptic vesicle, synaptic compartment, and active zone segmentation. We used the compartment segmentation to select only the vesicles of the presynaptic compartment of interest, by finding the compartment with most vesicles and then discarding all vesicles that did not overlap with it. We post-processed the active zone segmentation by intersecting the model predictions with a Sato filter^71^ that yields high responses for the active zone membrane, and skeletonizing the obtained mask to create a 2D surface of the active zone. We visually inspected the active zone segmentation results and found that 59 exhibited a good result. We proofread the other 42 tomograms, using painting in napari combined with a custom widget that snapped the painted area to the membrane (determined by the Sato edge filter) and skeletonized them. This process took about an hour and thirty minutes for all 42 tomograms. We measured the closest distance between synaptic vesicles and the active zone with an euclidean distance transform, scaled with the voxel size. The vesicle diameters were computed as twice the maxima of the euclidean distance to the vesicle boundary. We also derived the active zone surface area from a 3D mesh fitted to the segmented object via marching cubes. However, we found that this measurement was inconsistent compared to the area measurements obtained from IMOD, even when applied to the same segmented object. We thus did not analyze the normalized synaptic vesicle distances for the automatic segmentation results; for the corresponding results in Figures 4J and 5J we used the area of the manually segmented active zones for normalization. We used the implementations from SciPy^67^ and scikit-image^68^ for the image processing functions (image filters, skeletonization, euclidean distance transform, marching cubes).

For the inner ear ribbon synapses, a subset of tomograms was annotated for synaptic vesicles, vesicle pool identities, ribbon, presynaptic density, and active zone membrane in IMOD. For these annotations the synaptic vesicle annotations were extracted via the imodinfo command. Distances between synaptic vesicles and structures were measured by exporting the annotations from IMOD to segmentations and using an euclidean distance transform scaled with the voxel size of the data. SynapseNet was applied to all tomograms to segment the synaptic vesicles and structures. Distances and vesicle diameters were computed as for the other analysis (previous paragraph) and vesicle pools were assigned based on the distance criteria, see description in <u>Automatic Reconstruction and Analysis</u>. These results were proofread using napari in two different stages. First the structures (ribbon, presynaptic density and active zone membrane) were corrected via painting. Then the vesicle pool assignments were recomputed based on updated distances. In a second step, vesicle segmentations were corrected, e.g. to paint in missing vesicles or split wrongly merged vesicles, and vesicle pool assignments were corrected in cases where the assignments based on the distance measurements disagreed with manual judgment.

### Statistics

Statistical analysis was performed on Mathworks Graphpad Prism (versions 9.0 and 10.0). Samples were tested for normality distribution. For comparisons between two groups, the two groups were analyzed using Student’s unpaired t-tests and Mann-Whitney unpaired tests for normally and non-normally distributed data sets, respectively. For comparisons of multiple conditions (i.e. frequency distribution of SVs), statistical significance was tested by two-way ANOVA, with Bonferroni correction for multiple comparisons. Significant values from ANOVA tests were retested using either Student’s t-test for normally distributed data, and Mann-Whitney test for non-normally distributed data. The number of samples and biological replications used for each experiment are indicated in the corresponding figure legends.

## Code Availability

The SynapseNet software and all additional scripts for data analysis in this manuscript are available at https://github.com/computational-cell-analytics/synapse-net. The segmentation models can be downloaded with our software. We will submit them to BioImage.IO^72^, a community resource for sharing deep learning models for microscopy, upon acceptance of the manuscript.

## Data Availability

We use different, already published and unpublished, electron microscopy datasets in this manuscript, see <u>Sample Preparation and Data Preparation</u> and Supplementary Table 1 for an overview. We have made the data that was derived from published electron micrographs available via Zenodo. Data derived from yet unpublished micrographs will be made available on Zenodo after publication of the corresponding biological study. Until then, it is available upon request, on the condition that this data will only be used for developing or comparing segmentation methodology and not for biological analysis. Specifically, the datasets are available as follows:

- **Chemical Fixation** & **Single-Axis TEM Tomo**: Available on Zenodo: <u>10.5281/zenodo.14236426</u>
- **Dual-Axis TEM Tomo** & **STEM Tomo**: Will be made freely available after the corresponding publication. Available upon request.
- **IER**: The already published tomograms are available on Zenodo: <u>10.5281/zenodo.14232606</u>. The not yet published tomograms will be made freely available after the corresponding publication. Available upon request.
- **EH**: Will be made freely available after the corresponding publication. Available upon request.
- **Cryo**: Will be made freely available after the corresponding publication. Available upon request.
- **Frog**: Available on Zenodo: <u>10.5281/zenodo.14232529</u>
- **2D TEM**: Available on Zenodo: <u>10.5281/zenodo.14236381</u>
- **MUNC/SNAP**: Available on Zenodo: <u>10.5281/zenodo.14254111</u>

## Acknowledgments

This work was supported by the German Research Foundation (Deutsche Forschungsgemeinschaft, DFG) through grants SFB1286/A01 (to B.H.C., funded F.M. and I.H.-G.-P.), SFB1286/A03 (to S.O.R.), SFB1286/A04 (to C.W., funded J.N.B.), SFB1286/A12 (to R.F.-B., funded A.P.), SFB1286/B05/C08 (to T.M.), and SFB889/A07 (to C.W.). The DFG also supported the work of L.F., So.M. (Hertha Sponer College), C.W., R.F.-B., T.M., S.O.R., and C.P. under Germany’s Excellence Strategy - EXC 2067/1-390729940; supported A.A. through PA 4341/2-1; and supported N.B. through the NeuroNex International Network, BR 1107/15-2. C.P. was further supported by the Google Research Scholarship “Vision Foundation Models for Bioimage Segmentation” and T.M., Su.M. by the European Research Council through the Advanced Grant ‘DynaHear” under the European Union’s Horizon 2020 Research and Innovation program (grant agreement No. 101054467, TM). We also gratefully acknowledge the computing time granted by the Resource Allocation Board and provided on the supercomputer Lise and Emmy at NHR@ZIB and NHR@Göttingen as part of the NHR infrastructure. The calculations for this research were conducted with computing resources under the project nim00007. We thank S. Beuermann and the AGCT Lab for excellent technical support and are grateful to the MPINAT City Campus animal facility and precision mechanics workshop for their support throughout the project. We thank Rituparna Chakrabarti for providing three additional tomograms that were used for training segmentation models for ribbon synapse structure segmentation. She prepared these tomograms as a PostDoctoral researcher in the group of C.W.; the re-use of this data for this study started after her tenure. We thank Maksims Fiosins and Omar Diaz for supporting us with the set-up of IT infrastructure for this study.

## Author Contributions

F.M., So.M., I.H.-G.-P., J.N.B., A.P., T.T.D., V.S., K.W., Su.M., L.M., C.I., S.O.R., and B.H.C. acquired data. S.M., F.M., So.M., I.H.-G.-P., J.N.B., A.P., V.S., K.W., L.M., and C.P. annotated data for network training or evaluation. S.M., L.F., A.A., and C.P. implemented the software for network training, evaluation and data analysis in SynapseNet. S.M., F.M., So.M., B.H.C., and C.P. performed the data analysis for Figures 4-6. Su.M., N.B., C.W., R.F.-B., T.M., S.O.R., B.H.C., and C.P. conceptualized the work and supervised it. S.M., F.M., B.H.C., and C.P. drafted the manuscript. All co-authors read and revised the manuscript.

## Supplementary Tables

**Supplementary Table 1:** Summary of ultrastructural data used for supervised training and evaluation (yellow), domain adaptation (blue), and for application studies (red). Abbreviations: Data usage: Supervised training and evaluation, ST; domain adaptation, DA; Application study, AS. Synapse types and organelles: Schaffer collateral, SC; mossy fiber, MF; neuromuscular junction, NMJ; Endbulb of Held, EH; Inner ear ribbon synapses, IER; hippocampal cell culture, HCC; synaptic vesicles, SV; synaptic ribbons, SR; mitochondria, Mito. Fixation conditions: chemical fixation, CF; high-pressure freezing, HPF; automated freeze-substitution, AFS; plunge freezing, PF.

| EM Dataset |  | Sample Description |  |  | Sample Preparation / Image Acquisition |  | Data Usage |  |  |
| --- | --- | --- | --- | --- | --- | --- | --- | --- | --- |
| Name | Publication | Organism | Synapse | # | Fixation | Microscope | ST | DA | AS |
| Chemical Fixation | 15 | Mouse | SC/MF | 23 | CF/HPF/AFS | 200 kV JEM 2100 (JEOL) | SV/AZ<br>M |  |  |
| Single-Axis TEM Tomo | 12 | Mouse | SC | 152 | HPF/AFS | 200 kV JEM 2100 (JEOL) | SV |  |  |
| Dual-Axis TEM Tomo | unpublished | Mouse | HCC/MF | 114 | HPF/AFS | 200 kV Talos F200C (Thermo) | SV |  |  |
| STEM Tomo | unpublished | Mouse | SC/MF/PS | 48 | HPF/AFS | 200 kV Talos F200C (Thermo) | SV/AZ<br>M/SynC |  |  |
| IER | 11,38,39 and unpublished | Mouse<br>Rat | IER | 116 | HPF/AFS | 200 kV JEM 2100Plus (JEOL) | SR | SV | Fig. 6 |
| EH | unpublished | Mouse | EH | 44 | HPF/AFS | 200 kV JEM 2100Plus (JEOL) |  | SV |  |
| Cryo | unpublished | Rat | HCC | 22 | PF | 300 kV Krios G4 (Thermo) |  | SV |  |
| Frog | 43 | Frog | NMJ | 402 | CF | 80 kV CM10 (Philips) |  | SV |  |
| 2D TEM | 15 | Mouse | SC | 13 | HPF/AFS | 80 kV LEO 912 (Zeiss) |  | SV |  |
| Munc13 / SNAP-25 | 12 | Mouse | SC | 101 | HPF/AFS | 200 kV JEM 2100 (JEOL) |  |  | Fig. 4<br>Fig. 5 |

## Supplementary Figures

**Supplementary Figure 1:**
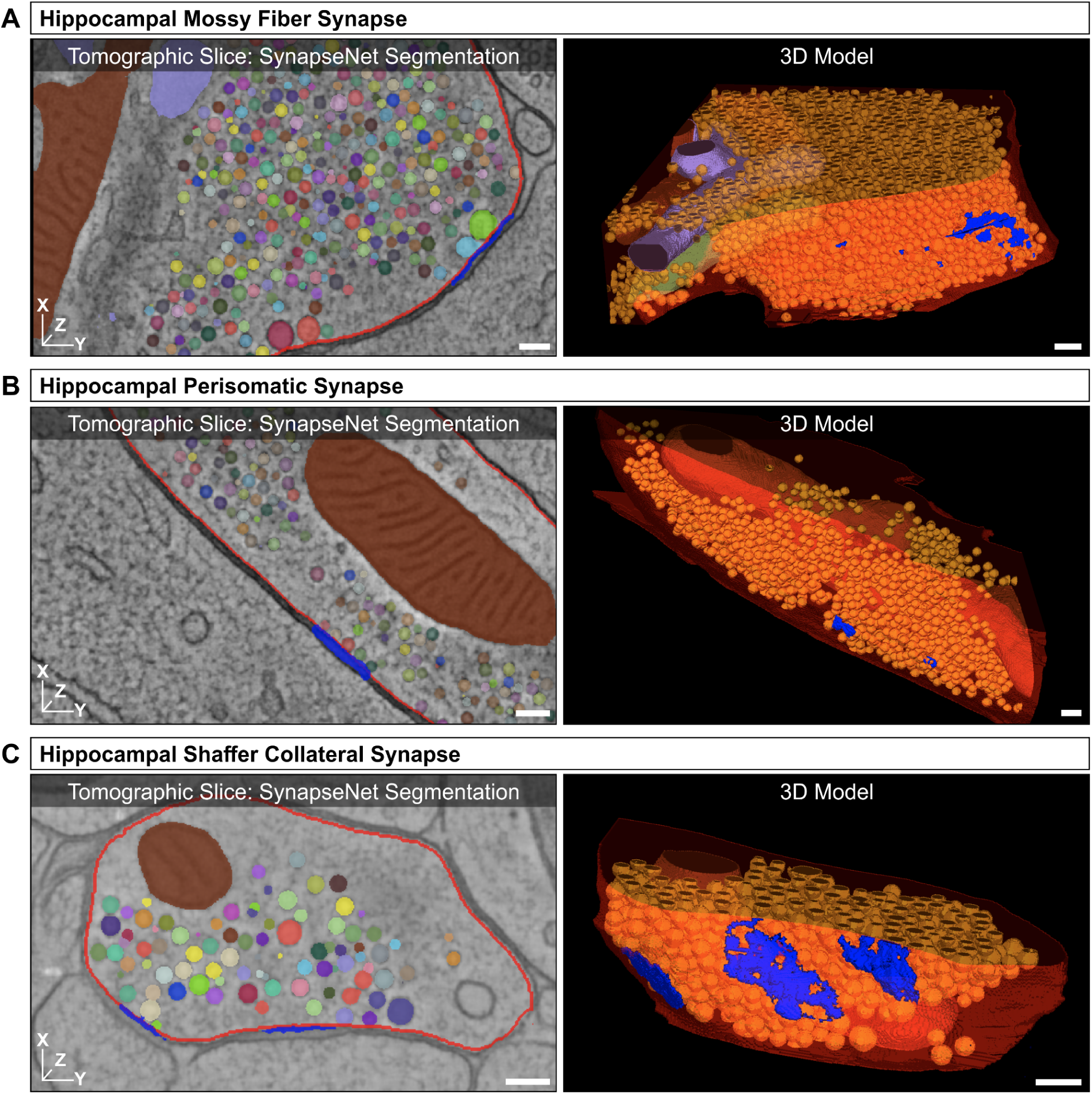
Automated synapse reconstructions of a mossy fiber (A), perisomatic (B) and Schaffer collateral (C) synapse from SynapseNet. Active zones are rendered in blue, mitochondria in dark red and purple, synaptic compartments in red, and synaptic vesicles in a different color per object in the virtual section (left) and in orange in the 3D rendering (right). To obtain the reconstructions, different pieces (2-5) of the synaptic compartment predictions were merged for a correct segmentation, active zones were manually enlarged in some areas, all other segmentations are shown as they were predicted by the model. The scale bars represent 100 nm.

**Supplementary Figure 2:**
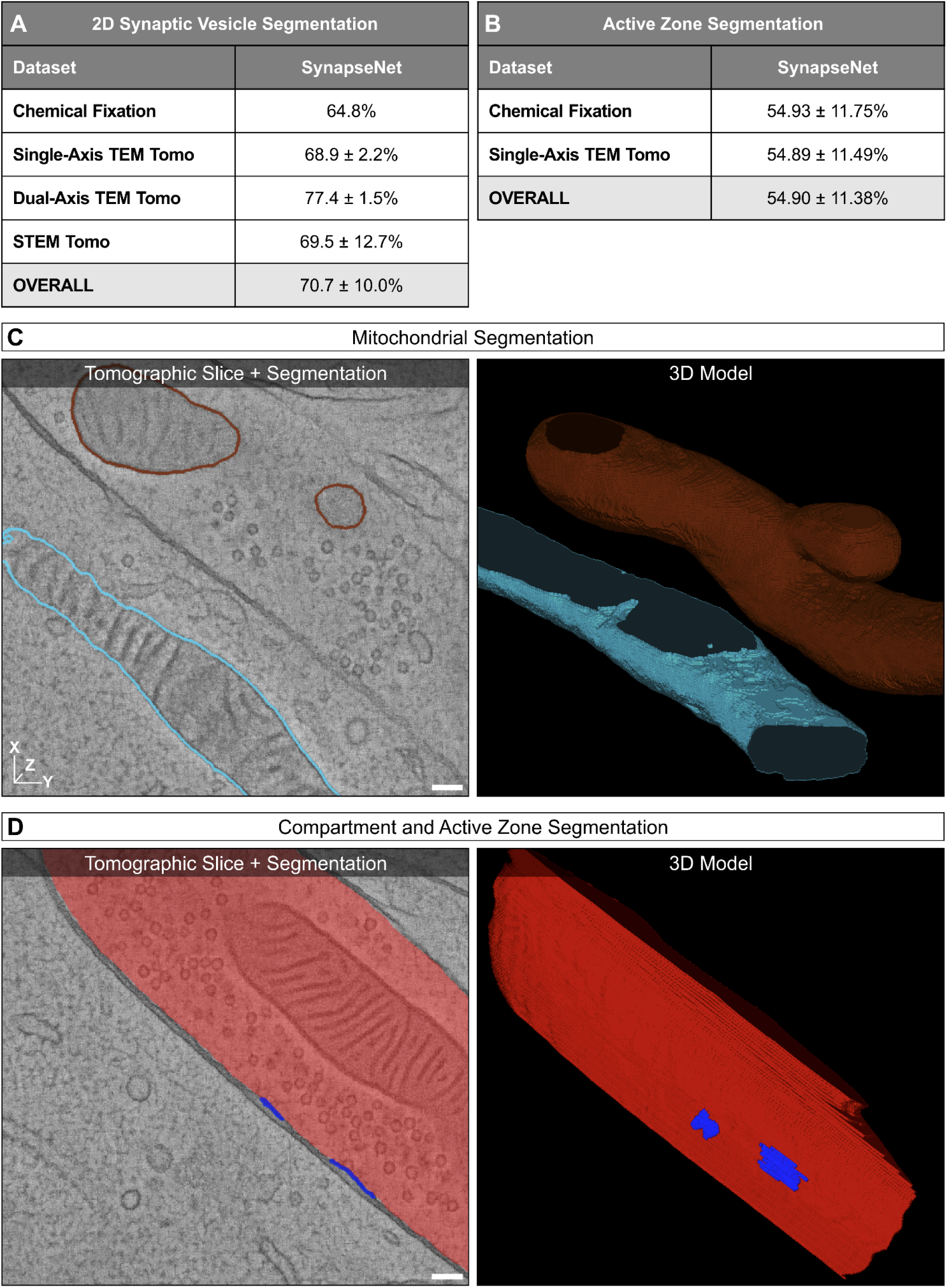
Segmentation evaluation and examples. **A**. Evaluation of 2D synaptic vesicle segmentation with SynapseNet, using the same set-up as in Figure 2C, except for segmentation evaluation on individual sections instead of in 3D. **B**. Evaluation of active zone segmentations. The Dice score is used for evaluation. **C**, **D.** Example segmentations for mitochondria, active zones, and a synaptic compartment in a perisomatic synapse, for a virtual section (left) and rendered in 3D (right). Mitochondria are rendered in and rendered in 3D in matching colors (right). **B**. Active zone (red) and synaptic compartment (blue) segmentations in a virtual section (left) and rendered in 3D. The segmentations are unedited. The scale bars represent 100 nm.

**Supplementary Figure 3:**
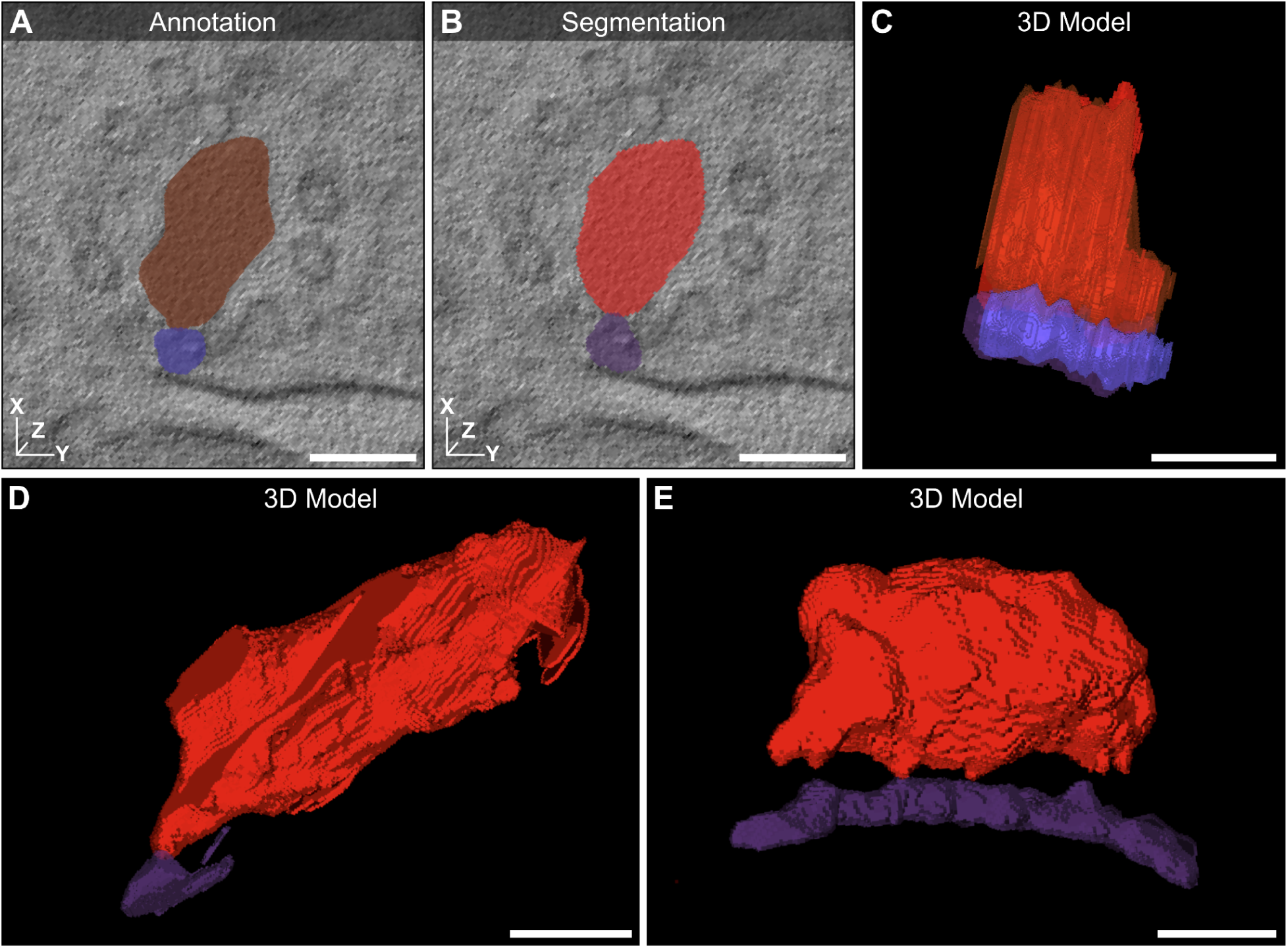
Example segmentations for ribbons and presynaptic densities in inner ear ribbon synapses. **A**,**B**. Annotation and SynapseNet segmentation of synaptic ribbon (red) and presynaptic density (purple) overlaid on top of a virtual section. **C.** 3D rendering of annotation and segmentation on top of each other. **D**,**E.** 3D renderings of two other ribbon and presynaptic segmentations. The scale bars represent 100 nm.

**Supplementary Figure 4:**
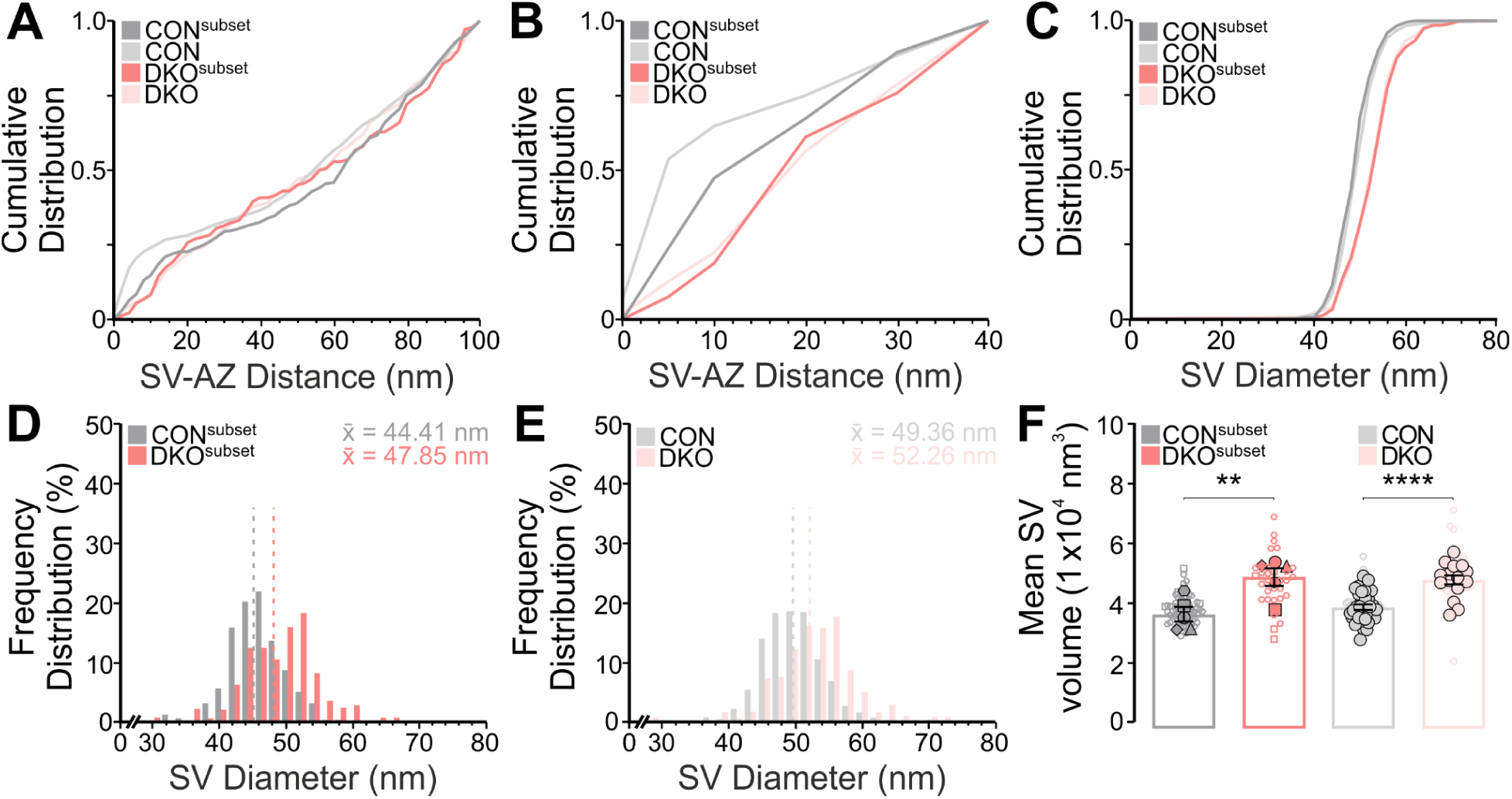
3D Electron tomographic analysis of synaptic vesicles in automatically segmented Munc13-1/2 double knock-out (DKO) and control (CON) hippocampal Schaffer collateral neurons. Analyses are based on automatic segmentation of synaptic vesicles (SVs) and semi-automatic segmentation of the active zone (AZ) from control and Munc13-1/2 DKO synapses. A subset of CON and 5 of DKO synapses (n=5) was compared with a larger dataset of electron tomographic subvolumes (CON, n=31; Munc13-1/2 DKO, n=15). **A**,**B**. Cumulative spatial distribution of SVs within 100 nm and 40 nm of the AZ. **C**. Cumulative distribution of SV diameters within 100 nm of the AZ. **D**,**E**. Frequency distribution of SV diameters within 100 nm of the AZ. Dotted lines indicate the mean SV diameter (x̅). **F**. Scatterplot of the mean volume of SVs within 100 nm of the AZ. Filled data points indicate the mean SV volume of single ET subvolumes and empty data points with the same symbol shape indicate the volume of single SVs within the same ET subvolume. Values indicate mean ± SEM; ****p < 0.0001; ***p < 0.001; **p < 0.01; *p < 0.05.

**Supplementary Figure 5:**
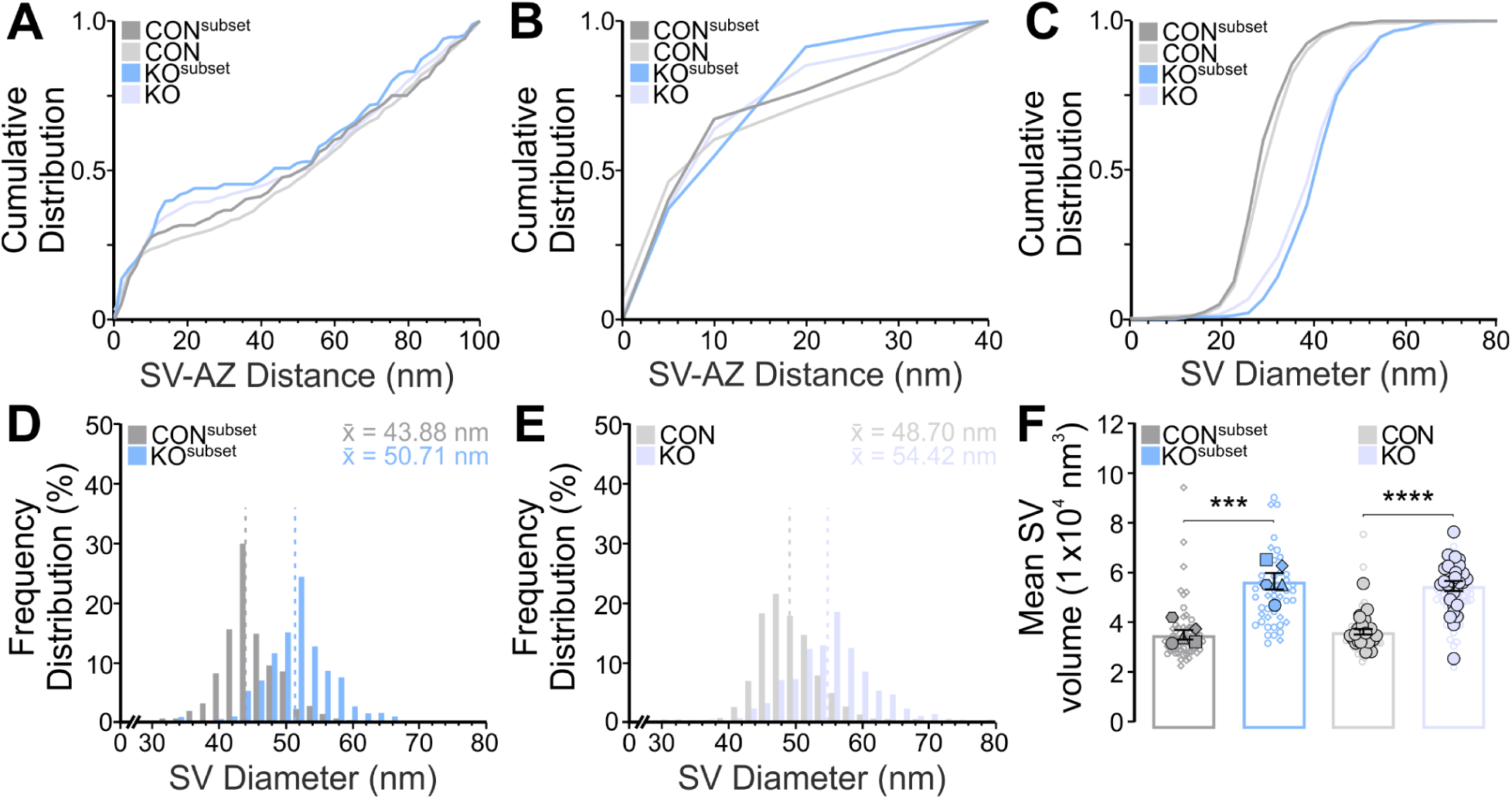
3D Electron tomographic analysis of synaptic vesicles in automatically segmented control (CON) and SNAP-25 knock-out (KO) hippocampal Schaffer collateral neurons. Analyses are based on automatic segmentation of synaptic vesicles (SVs) and semi-automatic segmentation of the active zone (AZ) from control and SNAP-25 KO synapses. A subset of CON (n=5) of KO synapses (n=5) was compared with a larger dataset of electron tomographic subvolumes (CON, n=27; SNAP KO, n=28). **A,B.** Cumulative spatial distribution of SVs within 100 nm and 40 nm of the AZ. **C.** Cumulative distribution of SV diameters within 100 nm of the AZ. **D,E.** Frequency distribution of SV diameters within 100 nm of the AZ. Dotted lines indicate the mean SV diameter (x̅). **F.** Scatterplot of the mean volume of SVs within 100 nm of the AZ. Filled data points indicate the mean SV volume of single ET subvolumes and empty data points with the same symbol shape indicate the volume of single SVs within the same ET subvolume. Values indicate mean ± SEM; ****p < 0.0001; ***p < 0.001; **p < 0.01; *p < 0.05.

**Supplementary Figure 6:**
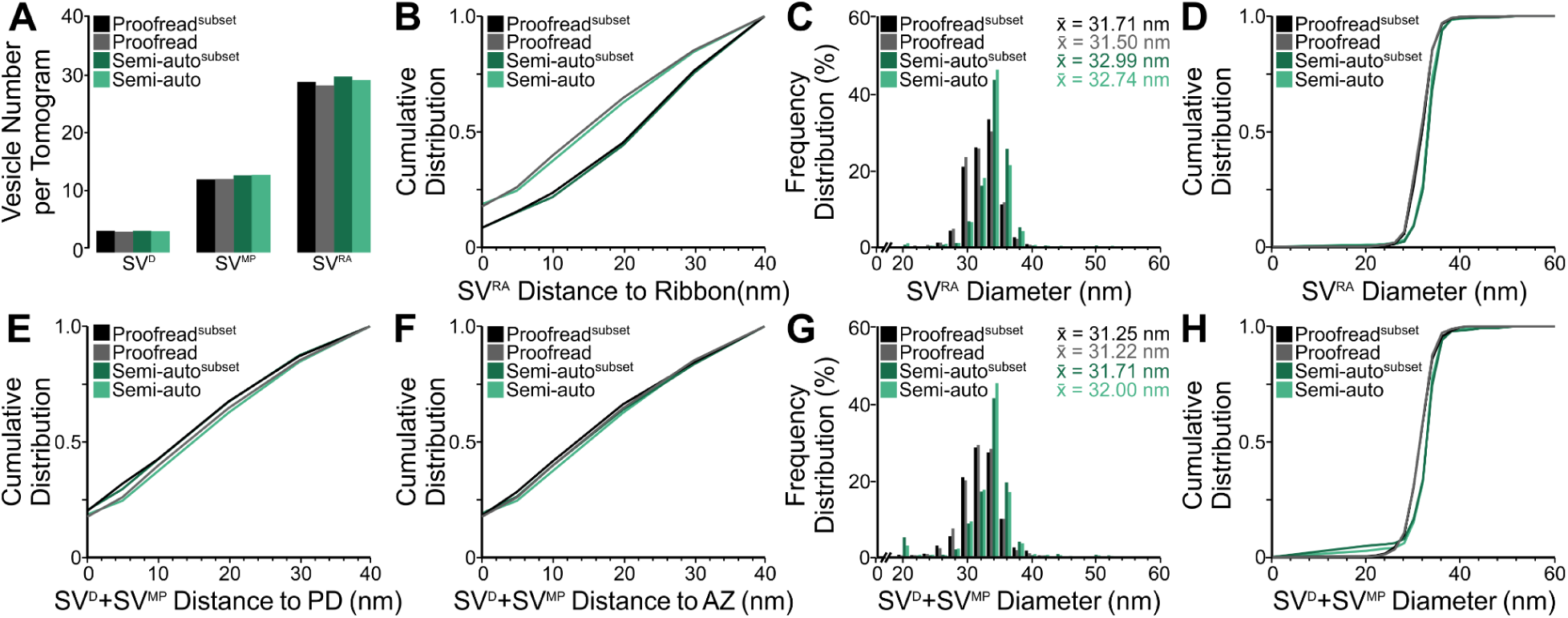
Analysis of ribbon synapses of cochlear inner hair cells in the inner ear. A subset (n=33) and a larger dataset of proofread and semi-automatic segmentations (n=88) were analysed. **A.** Average number of docked synaptic vesicles (SV^D^), membrane-proximal SVs (SV^MP^), and ribbon-associated SVs (SV^RA^) per tomogram. **B.** Cumulative spatial distribution of SV^RA^ within 40 nm of the ribbon. **C,D.** Frequency distribution of SV^RA^ diameters with the mean SV diameter (x̅) and cumulative distribution of SV diameters. **E,F.** Cumulative spatial distribution of SV^D^+SV^MP^ within 40 nm of the PD and the AZ. **G,H.** Frequency distribution of SV^D^+SV^MP^ diameters with the mean SV diameter (x̅) and cumulative distribution of SV diameters. Values indicate the mean diameter.

**Supplementary Figure 7:**
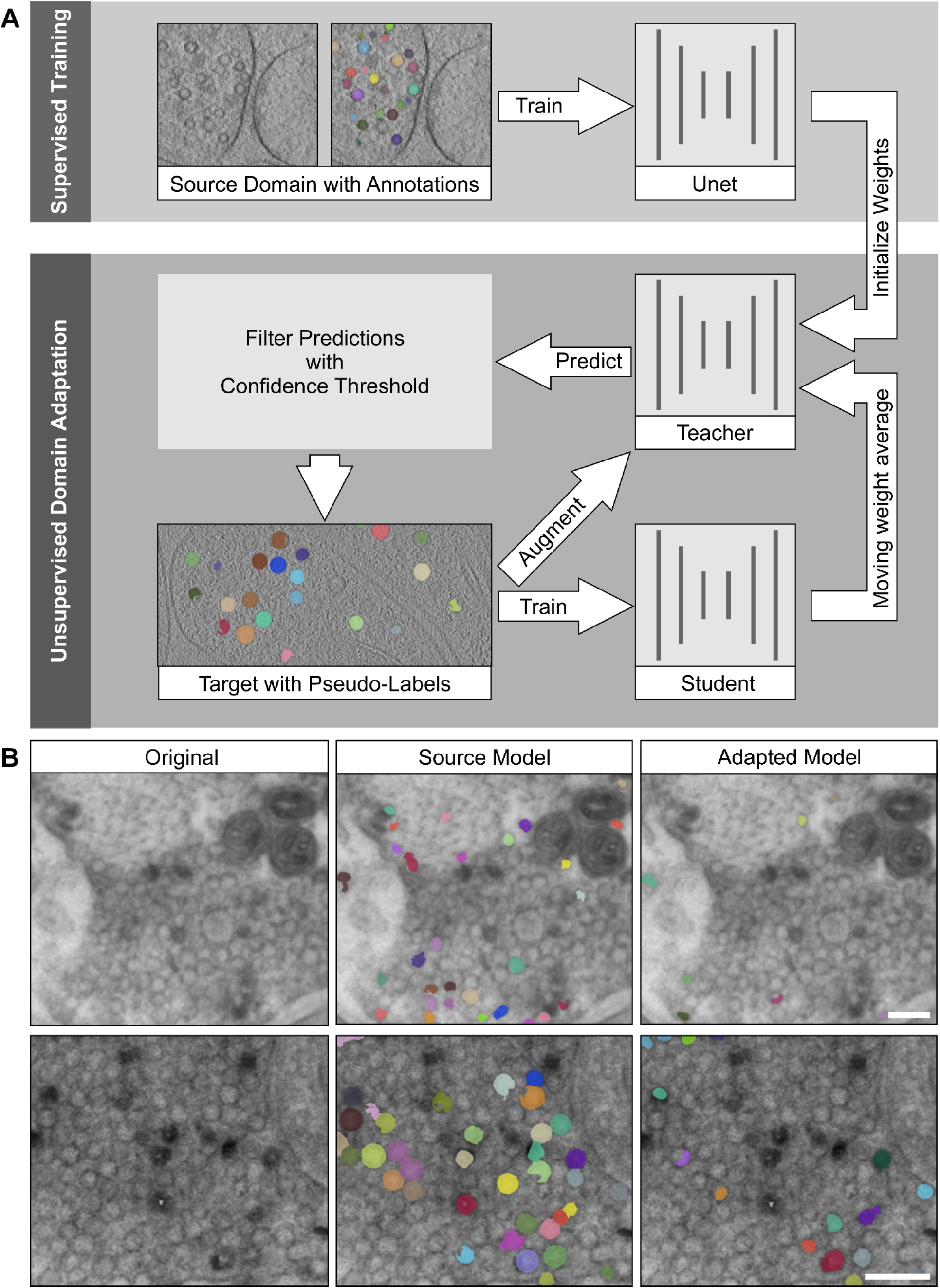
**A.** Schematic overview of domain adaptation. To transfer a model to a new domain (different imaging modality, sample preparation, specimen, etc.), we start from a model trained on the source domain, corresponding to the model trained via supervised training on data with annotations (top). For training on the target domain the model architecture is duplicated to obtain a teacher and a student model. In each training iteration the teacher model is applied to an augmented version of the image and its output is filtered to retain only confident predictions. The student model is applied to the image and its predictions are compared via a loss function to the filtered teacher output. The weights of the student network are updated via stochastic gradient descent (or a variant thereof) and the weights of the teacher network are updated via exponential moving average of the student network’s weights. **B.** Limitations of domain adaptation. If the initial predictions of the teacher network miss the majority of objects, e.g. vesicles, then the domain adaptation will likely fail and converge to predicting fewer or even no objects at all. This is the case for the frog data, which is shown here. The appearance of the image data is too different from the annotated vesicle training data due to lower contrast, leading to bad predictions of the source model and worse predictions after domain adaptation. The scale bars represent 200 nm.

